# Patient-Derived Mutant Forms of NFE2L2/NRF2 Drive Aggressive Murine Hepatoblastomas

**DOI:** 10.1101/2020.09.13.295311

**Authors:** Huabo Wang, Jie Lu, Jordan A. Mandel, Weiqi Zhang, Marie Schwalbe, Joanna Gorka, Ying Liu, Brady Marburger, Jinglin Wang, Sarangarajan Ranganathan, Edward V. Prochownik

## Abstract

**Background and Aims:** Hepatoblastoma (HB), the most common pediatric liver cancer, often bears β-catenin mutations and deregulates the Hippo tumor suppressor pathway. Murine HBs can be generated by co-expressing β-catenin mutants and the constitutively active Hippo effector YAP^S127A^. Some HBs and other cancers also express mutants of NFE2L2/NRF2 (NFE2L2), a transcription factor that tempers oxidative and electrophilic stress. In doing so, NFE2L2 either suppresses or facilitates tumorigenesis.

**Methods:** We evaluated NFE2L2’s role in HB pathogenesis by co-expressing all combinations of mutant β-catenin, YAP^S127A^ and the patient-derived NFE2L2 mutants L30P and R34P in murine livers. We evaluated growth, biochemical and metabolic profiles and transcriptomes of the ensuing tumors.

**Results:** In association with β-catenin+YAP^S127A^, L30P and R34P markedly accelerated HB growth and generated widespread cyst formation and necrosis, which are otherwise uncommon features. Surprisingly, any two members of the mutant β-catenin-YAP^S127A^-L30P/R34P triad were tumorigenic, thus directly establishing NFE2L2’s oncogenicity. Each tumor group displayed distinct features but shared 22 similarly deregulated transcripts, 10 of which perfectly correlated with survival in human HBs and 17 of which correlated with survival in multiple adult cancers. One highly up-regulated transcript encoded serpin E1, a serine protease inhibitor that regulates fibrinolysis, growth and extracellular matrix. The combination of mutant β-catenin, YAP^S127A^ and Serpin E1, while not accelerating cystogenic tumor growth, did promote the wide-spread necrosis associated with mutant β-catenin-YAP^S127A^-L30P/R34P tumors.

**Conclusions:** Our findings establish the direct oncogenicity of NFE2L2 mutants and key transcripts, including serpin E1, that drive specific HB features.

## Introduction

Hepatoblastoma (HB), the most common pediatric liver cancer, usually arises before three years of age. Factors impacting survival include age, α-fetoprotein levels, histologic subtype and transcriptional profiles ^1^. Heterogeneous mutations in the β-catenin transcription factor occur in ~80% of HBs in association with β-catenin’s nuclear accumulation ^2^. Most HBs also deregulate the Hippo pathway and also aberrantly accumulate its terminal effector YAP (yes-associated protein) in the nucleus ^2, 3^.

Mutations impair β-catenin’s interaction with the adenomatous polyposis coli complex that normally phosphorylates β-catenin and licenses its proteasome-mediated degradation ^1, 3, 4^. Stabilized β-catenin then enter the nucleus, associates the Tcf/Lef family of transcriptional co-regulators, up-regulates oncogenic drivers such as c-Myc and Cyclin D1 and initiates tumorigenesis ^1, 3, 4^. Hydrodynamic tail vein injection (HDTVI) of Sleeping Beauty (SB) plasmids encoding a patient-derived 90 bp in-frame N-terminal deletion of β-catenin [Δ(90)] and YAP^S127A^, a nuclearly-localized YAP mutant, efficiently promotes HB tumorigenesis whereas neither individual factor is tumorigenic ^4^.

Different β-catenin mutants uniquely influence HB features ^3^. For example, mice with Δ(90)- driven HBs survive ~11-13 wks. Their tumors display the “crowded fetal” histology of the most common human HB subtype and differentially express ~5300 transcripts relative to liver ^3, 4^. In contrast, tumors generated by the Δ(36-53) mutant grow slower, demonstrate greater histologic heterogeneity and differentially express >6400 transcripts ^3^. The causes of these differences are complex and determined by each mutant’s stability, nuclear localization and transcriptional potency ^3^.

HBs are less genetically complex than other cancers^5,6^. Nevertheless, ~5-10% of HBs also harbor recurrent missense mutations in the *NFE2L2/NRF2* (NFE2L2) gene and up to 50% have copy number increases^7^. Similar changes occur in adult cancers and correlate with shorter survival^8^. HBs with *NFE2L2* mutations are associated with shorter survival, although how β-catenin heterogeneity affects this is unknown^7^. Our profiling of 45 murine HBs generated by eight patient-derived β-catenin mutants identified one *NFE2L2* point mutation (L30P)^3^. These findings present largely anecdotal reckonings of NFE2L2’s role in HB, which cannot be further evaluated due to the small case numbers and β-catenin’s functional heterogeneity.

NFE2L2, a “Cap ’n’ Collar” bZIP transcription factor, mediates adaptations to oxidative, electrophilic and xenobiotic stresses^9,10^. NFE2L2 normally forms a cytoplasmic complex with Kelch-like ECH-associated protein 1 (KEAP1) via short NFE2L2 segments known as the ETGE and DLG domains. NFE2L2-KEAP1 complexes interact with Cullin 3, an E3 ubiquitin ligase that targets NFE2L2’s proteosomal degradation, and ensures low basal expression^11^. The afore-mentioned stresses prompt the oxidation of multiple KEAP1 cysteine thiol groups that maintain complex integrity^10,11^. Dissociated and stabilized NFE2L2 thus translocates to the nucleus, heterodimerizes with members of the Maf family of bZIP transcription factors and binds to anti-oxidant response elements (AREs) in numerous target genes. Their products restore redox balance, metabolize xenobiotics, counter stress and apoptosis, regulate metabolic pathways and maintain genomic and mitochondrial integrity^9–12^. Most NFE2L2 mutations reside within or near the ETGE or DLG domains and abrogate KEAP1 association thereby leading to constitutive NFE2L2 nuclear translocation^12,13^. Alternatively, copy number variants (CNVs) in *NFE2L2* and *KEAP1* create stoichiometric imbalance and allow nuclear accumulation of wild-type (WT) NFE2L2.

NFE2L2’s suppresses reactive oxygen species (ROS)- and electrophile-mediated genotoxicity in cancer susceptible *nfe2l2-/-* mice^14^. ROS-scavenging NFE2L2 target gene products include peroxiredoxins, thioredoxin and thioredoxin reductase whereas others detoxify and/or promote the excretion of α,β–unsaturated carbonyl-, epoxide- and quinone-containing moieties^15,16^. In contrast, NFE2L2 deregulation may facilitate tumor growth by increasing oxidative stress tolerance and allowing previously unattainable levels of oncogene-stimulated proliferation ^17, 18^. NFE2L2 target gene products unrelated to redox regulation also benefit tumor growth, angiogenesis and metastasis^19,20^.

We show here that two patient-derived NFE2L2 missense mutants, L30P and R34P, dramatically accelerate hepatic tumorigenesis by β-catenin mutants and YAP^S127A^. The tumors also possessed large necrotic areas and innumerable fluid-filled cysts, an otherwise rare HB feature^21^. The exceedingly rapid growth of β-catenin+YAP^S127A^+L30P/R34P tumors was associated with a more robust anti-oxidant response. When co-expressed with either Δ(90) or YAP^S127A^ individually, L30P and R34P were also transforming, thus establishing them as actual oncoproteins. HBs expressing each combination of Δ(90), YAP^S127A^ and L30P/R34P shared a “core” set of 22 similarly regulated transcripts whose expression correlated with long-term survival in HBs and other cancers. While not affecting tumor growth or cyst formation when co-expressed with Δ(90) and YAP^S127A^, one transcript, encoding the serine protease inhibitor serpin E1, generated highly necrotic tumors, thereby supporting the idea that these 22 common transcripts cooperatively contribute to different tumor phenotypes.

## Materials and Methods

More detailed Methods are provided in the Supplementary Files.

### Plasmid DNAs

SB vectors encoding human β-catenin mutants Δ(90) and R582W, YAP^S127A^, WT-NFE2L2, NFE2L2^L30P^ (L30P), NFE2L2^R34P^ (R34P), cyto-roGFP, mito-roGFP and Myc-epitope-tagged murine Serpin E1^3,22,23^ were purified with Plasmid Plus Midi columns (Qiagen, Inc., Germantown, MD) and administered to 6–8-week-old FVB mice via HDTVI (10 μg each) along with 2 μg of a non-SB vector encoding SB transposase^3,4,23^.

### Tumor induction

Mice were euthanized when tumors achieved maximal permissible size. Fresh tissues were used for metabolic studies or were apportioned into small pieces, snap-frozen in liquid nitrogen and stored at −80C. All injections, monitoring procedures and routine care and husbandry procedures were approved by The University of Pittsburgh Department of Laboratory and Animal Resources and the Institutional Animal Care and Use Committee.

### Oxphos measurements

OCRs were determined with an Oroboros Oxygraph 2k instrument (Oroboros Instruments, Inc., Innsbruck, Austria) using partially purified mitochondrial suspensions in MiR05 buffer as previously described^3,23^.

### Oxidative stress

Isolated tumor cells maintained *in vitro* were exposed to 5 mM H_2_O_2_ followed by rapid replacement with H_2_O_2_-free medium. Monitoring of mito-rhoGFP and cyto-rhoGFP by live confocal microscopy was maintained over ~40 min to allow redox normalization rates to be determined.

### TaqMan Assays

To identify *NFE2L2* gene CNVs, DNAs from de-identified FFPE specimens of normal tissues and HBs were isolated using a QIAmp DNA FFPE Tissue Kit (Qiagen). All tissues were de-identified and were obtained with the approval of the University of Pittsburgh’s Institutional Review Board (IRB) guidelines and were in accordance with the Declaration of Helsinki principles. 10 ng of DNA was amplified in triplicate 12 μl reactions using a KAPA Probe Fast qPCR Kit (2x) (GeneWorks, Thebarton, Australia). mtDNA content was quantified using a previously-described TaqMan-based assay and normalized to the nuclear apolipoprotein B gene^3,24^. The nuclear genes glyceraldehyde-3-phosphate dehydrogenase (GAPDH) and Ribonuclease P RNA Component H1 were used for mtDNA normalization controls.

### Immuno-blotting

Immunoblotting and sub-cellular fractionation were performed as previously described using the antibodies and conditions shown in Supplementary Table S5^3,24^.

### RNAseq

RNA purification, RNAseq and bioinformatics processing and analyses were performed as previously described^3,24^.

### Target gene promoter analysis

Human and mouse genomes and gene promoters were accessed through the BSgenome.Hsapiens.UCSC.hg38 and BSgenome.Mmusculus.UCSC.mm10 R packages, respectively. Chip-Seq Data were accessed from the ENCODE v.5 database (https://www.encodeproject.org/).

### Dysregulation of NFE2L2 and its direct target genes in human cancers

RNAseq data from human HBs ^25,26^ were searched for NFE2L2 mutations using CLC Genomics Workbench 11. NFE2L2 and KEAP1 transcripts were downloaded from the TCGA PANCAN dataset as FPKM, converted to TPM, and filtered to contain only data from HCCs. CNVs across multiple cancers were identified by analyzing TCGA genomic data in Genomic Data Commons (GDC).

IPA was used to identify 46 promoters containing established NFE2L2-binding ARE elements. Expression data (TPM) for these genes and NFE2L2 itself were accessed from the GDC TCGA dataset available on the UCSC Xenabrowser (xena.ucsc.edu) from cohorts with the highest levels of NFE2L2 CNVs.

### Statistical analyses

Survival data for patients in TCGA, were analyzed with GraphPad Prism 7 (GraphPad Software, Inc., San Diego, CA).

## Results

### NFE2L2 mutants accelerate tumorigenesis

β-catenin mutations and WT β-catenin itself are tumorigenic when co-expressed with YAP^S127A 2–4^, with specific tumor features and transcriptomes determined by the β-catenin mutant’s identify^3^. Despite these differences, endogenous NFE2L2 transcripts were up-regulated approximately two-fold in all tumor groups relative to control livers (Supplementary Fig. S1A). Similar induction was observed in slower growing Δ(90)+YAP^S127A^ tumors lacking Myc, ChREBP or Myc+ChREBP ^23^. Like Myc, ChREBP is a transcription factor that regulates target genes involved in ribosomal biogenesis and carbohydrate and lipid metabolism^23^. These results indicate that, regardless of growth rates or other features, NFE2L2 is induced equally in murine HBs whereas KEAP1 expression is unchanged (Supplementary Fig. S1B).

*NFE2L2* missense mutations occur in 5-10% of HBs and at higher frequencies in other cancers^7,27,28^. To test the impact of these on survival, we injected SB vectors encoding Δ(90)+YAP^S127A^, with or without WT-NFE2L2, L30P or R34P. As expected, Δ(90)+YAP^S127A^ tumor-bearing mice had median survivals of ~11-13 wk (Fig. 1A)^4,24^, which were unaltered by co-injection with WT-NFE2L2. In contrast L30P and R34P co-expression significantly shortened survival (Fig. 1A).

**Figure 1.**
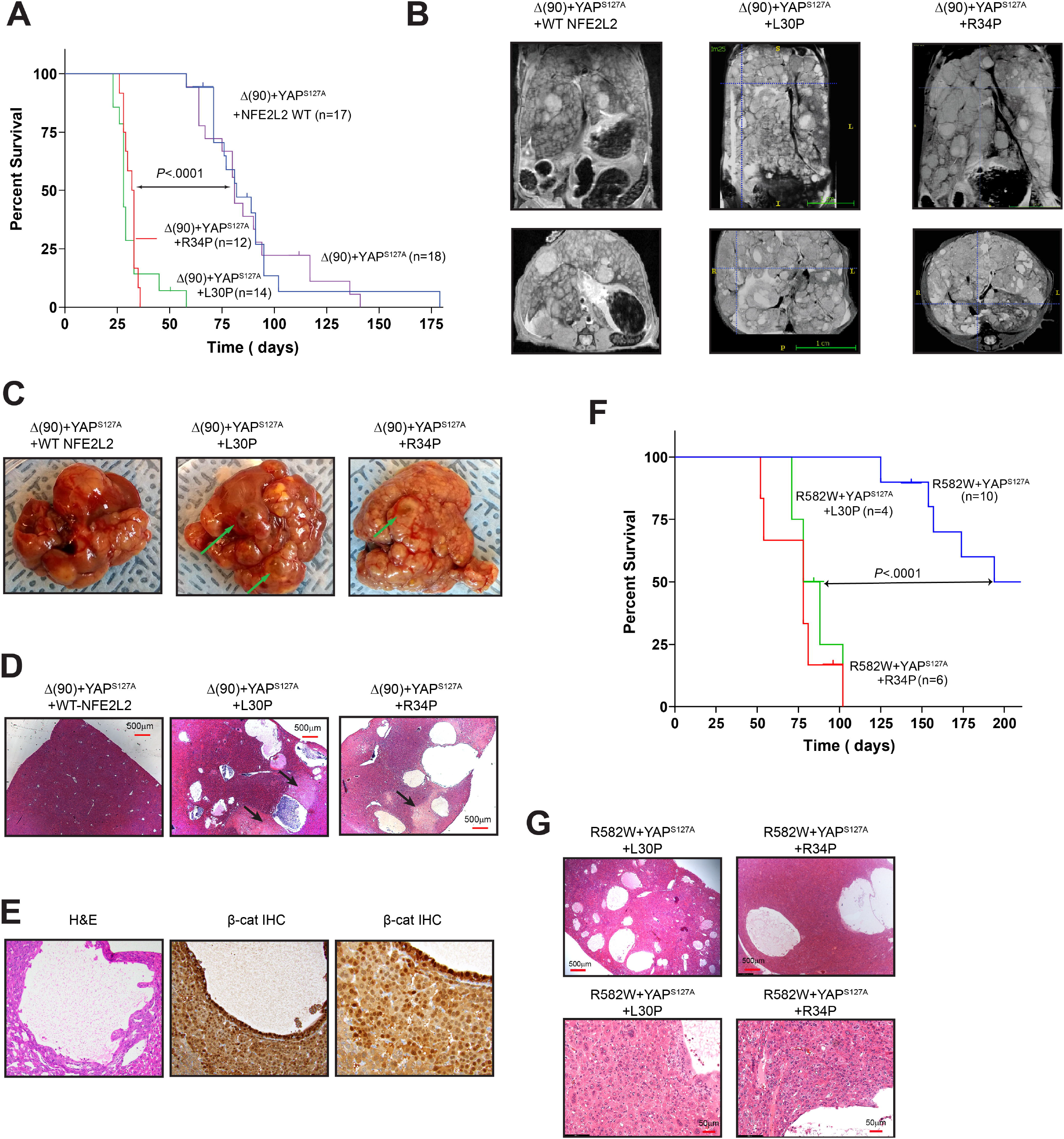
NFE2L2 mutants L30P and R34P accelerate HB growth. **A,** Kaplan-Meier survival curves of the indicated cohorts, n=10-12 mice/group. **B,** MRI images of comparably sized tumors just prior to sacrifice. **C,** Gross appearance of typical tumors from each of the groups with examples of typical fluid-filled cysts indicated by arrows. **D,** H&E-stained sections of the tumors shown in C showing multiple cysts with areas of prominent adjacent necrosis indicated by arrows. **E,** Higher power magnification of H&E- and β-catenin IHC-stained sections showing the lumens of cysts lined with cells resembling tumor cells that stain strongly for nuclearly-localized β-catenin. **F,** Kaplan-Meier survival curves of mice expressing β-catenin missense mutant R582W, YAP^S127A^ and the indicated NFE2L2 proteins. **G,** H&E stained sections of the indicated tumors from **F.**

L30P/R34P-expressing tumors displayed the crowded fetal pattern of Δ(90)+YAP^S127A^ HBs^4,24^. Additionally, they contained extensive areas of necrosis and innumerable fluid-filled cysts (Fig. 1B-D and Supplementary Fig S2). Cells lining the cysts were indistinguishable from tumor cells and stained intensely for nuclearly-localized β-catenin but not for the endothelial marker CD34 (Fig. 1E and data not shown) thus confirming their derivation from epithelium (Fig. 1E and Supplementary Fig. S2)^29^. Δ(90)+YAP^S127A^ tumors contained neither cysts nor significant necrosis.

Together with YAP^S127A^, the R582W β-catenin mutant generates slowly-growing tumors that more closely resemble hepatocellular carcinomas (HCCs)^3^. L30P and R34P also accelerated these tumors’ growth and generated numerous cysts (Fig. 1F&G and Supplementary Fig. S3).

### L30P and R34P are stabilized, nuclearly-localized and alter metabolic and redox states

L30P and R34P were modestly over-expressed relative to WT or endogenous NFE2L2 protein in livers and Δ(90)+YAP^S127A^ tumors and KEAP1 was expressed equally across cohorts (Fig. 2A and Supplementary Fig. S1B). Relative to control livers, tumor cohorts expressed distinct patterns of glucose transporters. For example, L30P/R34P tumors up-regulated ubiquitously-expressed Glut1, consistent with their rapid growth and greater reliance on Warburg-type respiration (Fig. 2A)^3,23,30^. They also tended to down-regulate liver-specific Glut2 and adipose- and muscle-specific Glut4. In contrast, Δ(90)+YAP^S127A^ tumors and Δ(90)+YAP^S127A^+WT-NFE2L2 tumors both up-regulated Glut4 and modestly down-regulated Glut2. All tumors markedly up-regulated the M2 isoform of pyruvate kinase (PKM2), which is favors tumor growth due to its lower K_m_ for phosphoenolpyruvate, leading to the accumulation of glycolytically-derived anabolic substrates^30^. Finally, all tumors expressed lower levels of carnitine palmitoyltransferase I (Cpt1a), β-FAO’s rate-limiting enzyme^31^. KEAP 1 was cytoplasmically localized (Fig. 2B) whereas WT NFE2L2, despite being cytoplasmic in most cell lines was nuclear in livers and HBs as was Δ(90). WT-NFE2L2 nuclear localization may reflect the liver’s heavy engagement in xenobiotic metabolism^10,12^.

**Figure 2.**
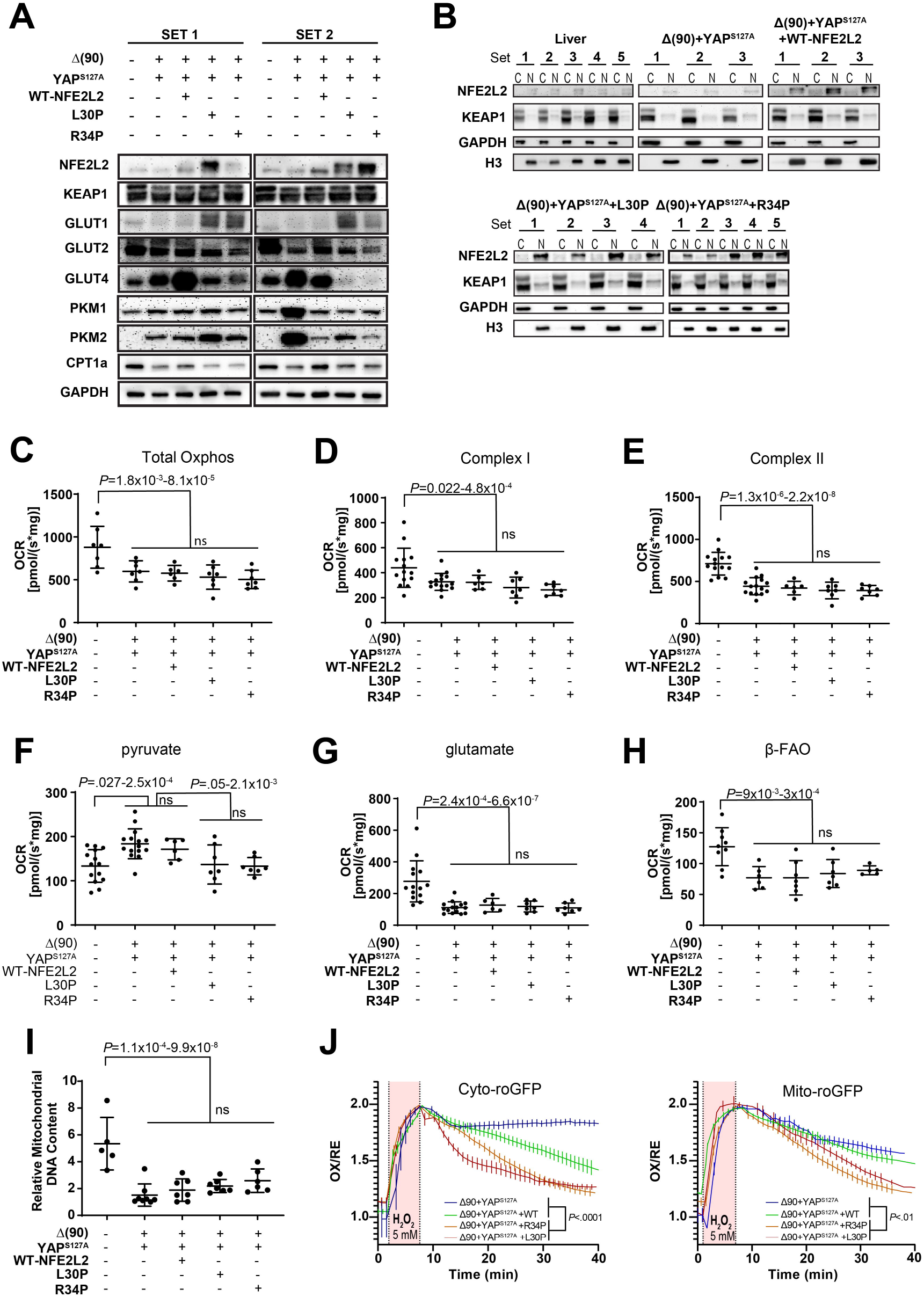
Distribution and metabolic consequences of L30P/R34P expression in HBs. **A,** Expression of NFE2L2, KEAP1, GLUT1, GLUT2, GLUT4, PKM1, PKM2 and Cpt1a in two representative sets of total lysates from the indicated tissues. **B,** Nuclear (N)/cytoplasmic (C) fractionation of the indicated tissues n=3-5 samples/group. GAPDH and histone H3 (H3) immunoblots were performed as controls for protein loading and the purity of each fraction. **C,** Total OCRs of mitochondria from the indicated tissues in the presence of malate, ADP, pyruvate, glutamate and succinate. **D,** Complex I responses, calculated following the addition of rotenone to the reactions in **C** without succinate. **E,** Complex II responses as determined from residual activity following the addition of Rotenone. **F,** Responses to pyruvate. **G,** Responses to glutamate. **H,**β-FAO responses following the addition of malatae, L-carnitine, and palmitoyl-CoA. **I,** Quantification of mitochondrial DNA (mtDNA) in representative tissues. TaqMan reactions amplified a segment of the mt D-loop region ^3, 24^. Each point represents the mean of triplicate TaqMan reactions after normalizing to a control TaqMan reaction for the ApoE nuclear gene. **J,** *In vitro* recovery from oxidative stress. Monolayer cultures of the indicated tumor cells expressing cyto-roGFP or mito-roGFP were exposed to 5 mM hydrogen peroxide (bar) while being monitored by live cell confocal microscope.

To assess L30P/R34P’s influence on metabolic functions, we examined mitochondrial oxygen consumption rates (OCRs) in response to TCA cycle substrates. Consistent with the switch from oxidative phosphorylation (Oxphos) to Warburg respiration, total OCRs and individual Complex I and Complex II contributions were reduced across all tumors groups relative to livers (Fig. 2C-E). Further consistent with previous observations^3,23,32^, all tumors increased their pyruvate response while suppressing their glutamate response (Fig. 2F and G).

HBs suppress β-FAO as they shift to glycolysis and hepatic lipid is mobilized for *de novo* membrane synthesis^3,23,32^. Regardless of NFE2L2 status, all tumors also down-regulated β-FAO (Fig. 2H), in keeping with lowever Cpt1a expression (Fig. 2A) and a reduction in mitochondrial DNA contnent as previously described (Fig. 2I) ^23^.

We next generated tumors expressing redox-sensitive forms of cytoplasmic- or mitochondrial-localized GFP (cyto-roGFP and mito-roGFP, respectively)^22^. Tumor cells were cultured for several days and briefly exposed to H_2_O_2_ while monitoring recovery by live-confocal microscopy. Δ(90)+YAP^S127^+WT-NFE2L2 cells recovered somewhat more rapidly than Δ(90)+YAP^S127A^ cells but this was significantly accelerated in tumor cells expressing L30P/R34P (Fig. 2J).

### L30P/R34P are tumorigenic in combination with eitherΔ(90) or YAP^S127A^

Rapid tumorigenesis mediated by L30P/R34P (Fig. 1A) raised questions regarding the latters’ role(s) in transformation. While no individual vector was oncogenic (Fig. 3A)^4,23^, L30P and R34P each generated tumors when co-expressed with Δ(90) or YAP^S127A^ with survivals being comparable to those of the R582W+YAP^S127A^ cohort (Figs. 1F, 3A)^4^. Thus, any pairwise combination of Δ(90), YAP^S127A^ and L30P/R34P is oncogenic. Δ(90)+L30P/R34P tumors were highly differentiated whereas YAP^S127A^+L30P/R34P tumors resembled HCCs with HB-like features (Fig. 3B, and Supplementary Fig. S4)^3,24^.

**Figure 3.**
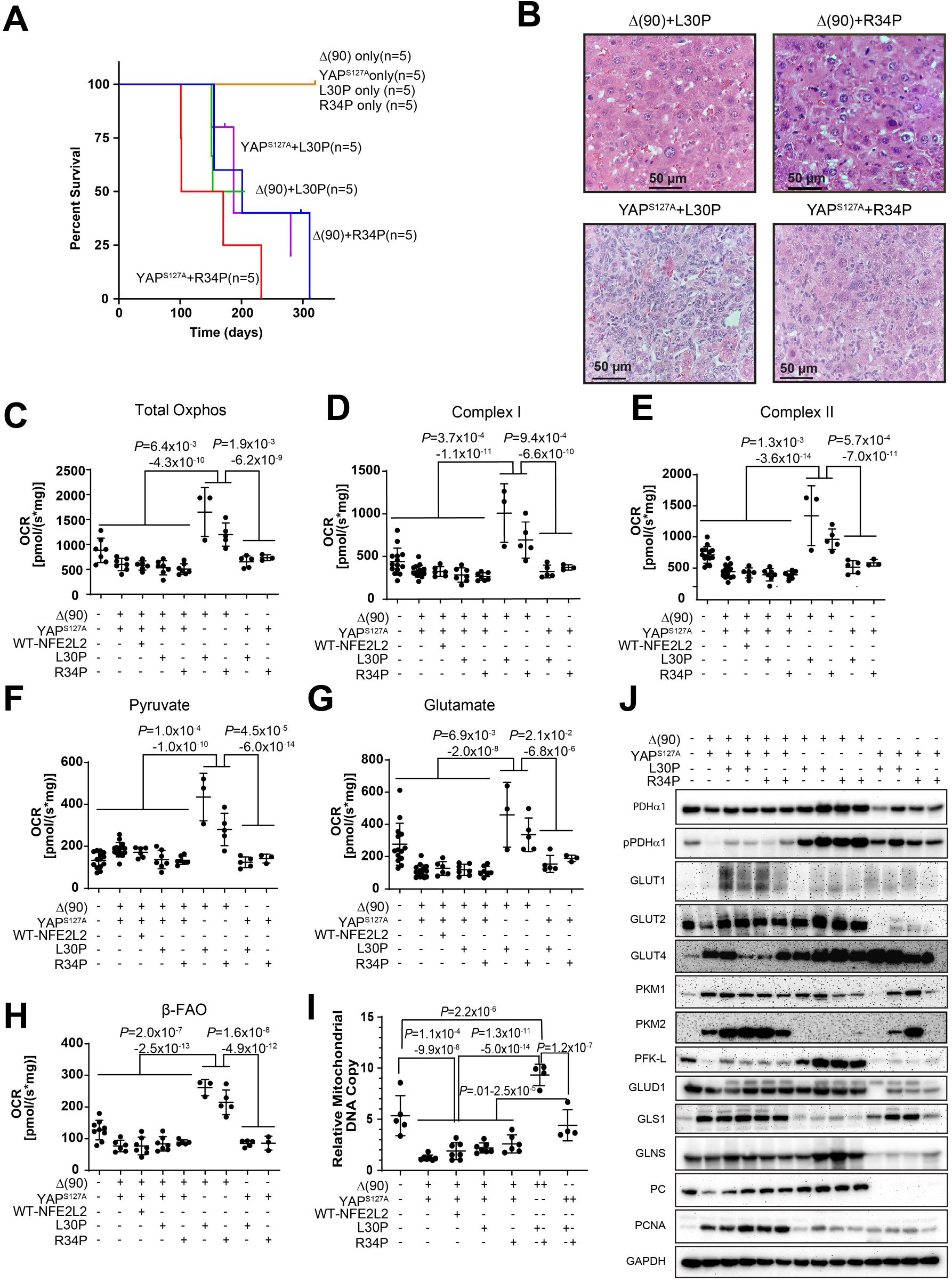
Characteristics of tumors generated by L30P and R34P co-expressed with Δ(90) or YAP^S127A^. **A,** Kaplan-Meier survival curves. Survival was determined as described in Fig. 1A. **B,** Histopathologic features of representative tumors from the indicated cohorts. See Supplementary Fig. S4 for additional images. **C-H,** OCRs performed as described in Fig. 2 **C-H. C**=Total Oxphos; **D**=Complex I; **E**=Complex II; **F**=pyruvate response; **G**=glutamate response; **H**= β-FAO. **I,** mtDNA content of representative tumors from the indicated cohorts performed as described in Fig. 2I. **J,** Immunoblots from representative tissues of the indicated cohorts. GAPDH was used as a loading control.

Metabolically, YAP^S127A^+L30P/R34P tumors were similar to those previously described (Fig. 3C-E) ^3, 24^ whereas Δ(90)+L30P/R34P tumors were distinct, with higher total OCRs and Complex I and II activities. Rather than the reciprocal relationship between β-FAO and pyruvate consumption and lower glutamate consumption documented previously (Fig. 2F&H)^3,24^ all activities, and mtDNA content, were increased in Δ(90)+L30P/R34P and exceeded even those of livers (Fig. 3F-I).

Components of the above pathways were again expressed in complex and cohort-specific ways. For example, pyruvate dehydrogenase α1 catalytic subunit (PDHα1) expression generally reflected each cohort’s mtDNA content (Fig. 3J). However, Ser_293_-phosphorylated PDHα1 (pPDHα1), was lower in Δ(90)+YAP^S127A^ tumors, irrespective of L30P/R34P status, indicating PDH activation and likely explaining the higher OCRs in response to pyruvate (Fig. 3F)^3,25,30^. Most notable were the higher pPDHα1 levels in Δ(90)+L30P/R34P and YAP^S127A^+L30P/R34P tumors, whose relative levels of PDH inactivation were similar (Supplementary Fig. S5).

Compared to Δ(90)+YAP^S127A^ and Δ(90)+YAP^S127A^+L30P/R34P tumors, Δ(90)+L30P/R34P and YAP^S127A^+L30P/R34P tumors down-regulated Glut1 and up-regulated Glut4. Most notable however was the marked down-regulation of Glut2 expression by the latter cohort (Fig. 2A and Fig. 3J). Δ(90)+L30P/R34P and YAP^S127A^+L30P/R34P tumors also tended to down-regulate PKM2, suggesting that it was not necessarily associated with transformation but rather reflected rapid growth rates.

The higher pyruvate utilization and greater mitochondrial activity of Δ(90)+L30P/R34P tumors were further accompanied by increased expression of the rate-limiting liver-type phosphofructokinase (PFK-L) (Fig. 3J). This, together with reduced PKM-2 expression, suggested that the more robust Oxphos of these tumors (Fig. 3C-H) was associated with increased glycolytic flow into the TCA cycle rather than into anabolic pathways that support rapid growth. Pyruvate production and its use for Oxphos in these different cohorts were thus under complex but cooperative control by factors that coordinated glucose uptake and its fate.

The complex Oxphos dependencies were further underscored by the glutamate response, which was reduced in most tumor groups relative to liver but up-regulated in Δ(90)+L30P/R34P tumors (Fig. 3G). Yet, levels of mitochondrial glutamate dehydrogenase 1 (Gludh1), which catalyzes glutamate’s conversion to α-ketoglutarate, were unchanged and those of mitochondrial glutaminase 1 (Gls1), which catalyzes glutamine conversion to glutamate, were actually reduced (Fig. 3J). Levels of glutamine synthase (Glns), which converts cytoplasmic glutamate to glutamine, were somewhat higher, suggesting that intratumoral glutamine availability poses a barrier to more extensive glutaminolysis. The extremely low levels of Gludh1 in YAP^S127A^+L30P/R34P tumors, which might limit the supply of α-ketoglutarate, were accompanied by a parallel marked decline of pyruvate carboxylase (PC), which anaplerotically furnishes oxaloacetate from pyruvate. As already demonstrated, the expected increase in pyruvate availability was not associated with greater pyruvate utilization. Instead, pyruvate might now prove a more abundant source of α-ketoglutarate via alanine transaminase. Finally, proliferating cell nuclear antigen (PCNA) levels closely matched tumor growth rates and survival (Figs. 1A&3A).

RNAseq analysis on the above cohorts demonstrated that most tumors reduced expression of transcripts encoding components of the TCA cycle, β-FAO, the ETC and ribosomes (Supplementary Fig. S6). The sole exception was seen in Δ(90)+L30P/R34P HBs which up-regulated these pathways as expected.

### RNAseq reveals common transformation-specific transcripts

The finding that any two members of the Δ(90)+YAP^S127A^+L30P/R34P trio were oncogenic (Figs. 1A&3A) suggested a model of tumorigenesis (and perhaps cystogenesis and necrosis), involving common transcripts that we sought to identify by RNAseq. Principal component analysis (PCA) and hierarchical clustering readily distinguished tumor cohorts from from liver (Fig. 4A and B). Because Δ(90)+YAP^S127A^+L30P and Δ(90)+YAP^S127A^+R34P tumors differed in the expression of only one transcript, they were subsequently combined. All other cohorts were distinguishable (Fig. 4B and C). Δ(90)+YAP^S127A^ tumors and Δ(90)+YAP^S127A^+WT-NFE2L2 tumors differed in the expression of 821 genes (Fig. 4C&D), indicating that WT-NFE2L2 was not entirely silent as suspected from its nuclear localization and modest redox buffering (Fig. 2B and J). Δ(90)+YAP^S127A^+L30P/R34P tumors showed more extensive transcriptional dysregulation, differing from Δ(90)+YAP^S127A^ and Δ(90)+YAP^S127A^+WT-NFE2L2 tumors by 3584 transcripts and 1654 transcripts, respectively (Fig. 4D).

**Figure 4.**
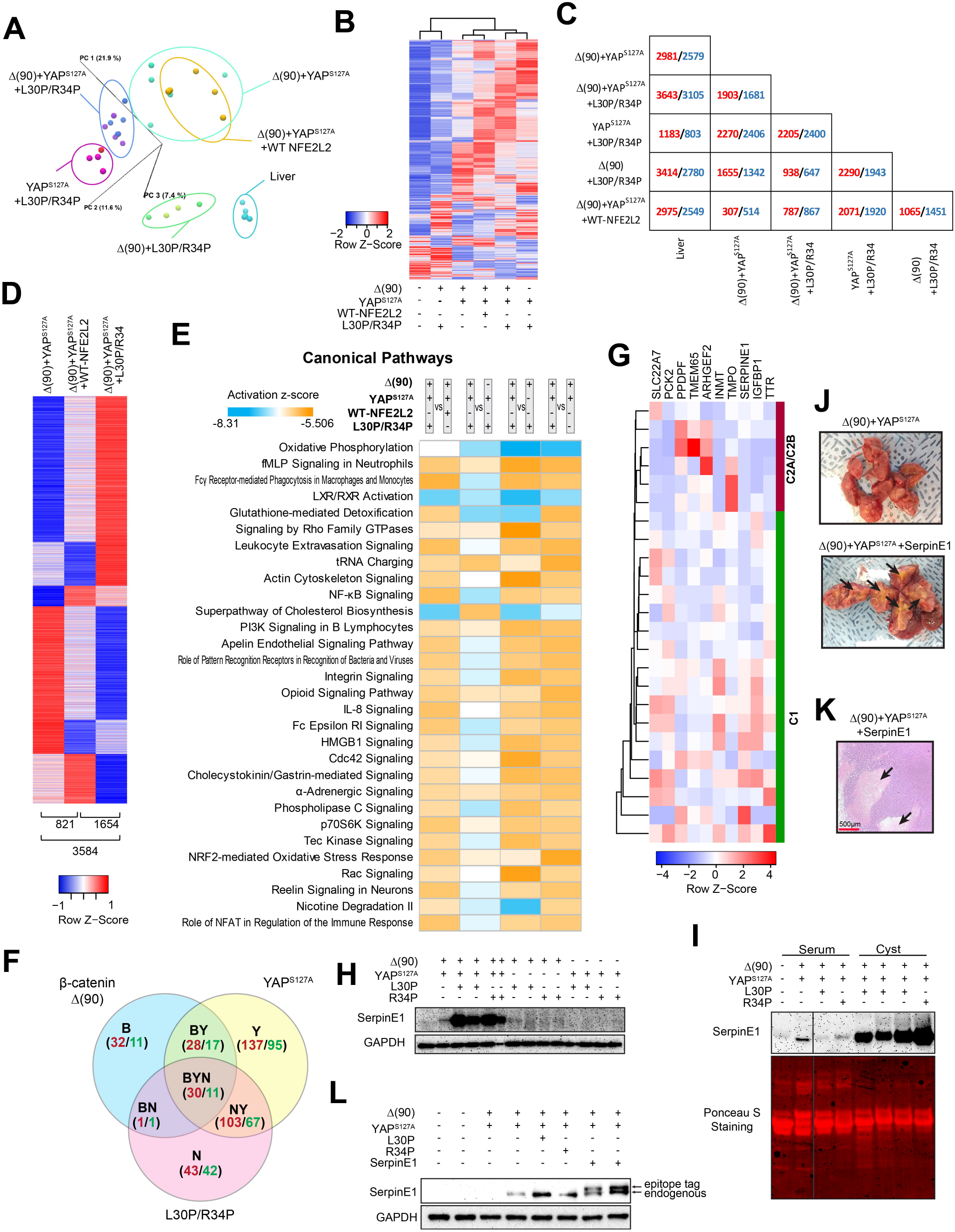
RNAseq analysis of tumors generated by combinations of Δ(90), YAP^S127A^, WT-NFE2L2, L30P and R34P. **A,** PCA of transcriptomic profiles of livers and tumors (n=4-5 samples/group. **B,** Heat maps of differentially expressed transcripts from the tissues depicted in (A) arranged by hierarchical clustering. Because only a single transcript difference was found between tumors expressing L30P and R34P, results were combined for this and subsequent analyses (L30P/R34P). Red=up-regulated in tumor vs. liver; blue=down-regulated in tumor vs. liver. **C,** Pairwise comparisons showing the number of significant gene expression differences between any two of the tissues depicted in **B.** Red and blue=up- and down-regulated, respectively, in the tumors depicted at the left relative to those indicated at the bottom. **D,** Distinct transcript patterns of Δ(90)+YAP^S127A^, Δ(90)+YAP^S127A^+WT-NFE2L2 and Δ(90)+YAP^S127A^+L30P/R34P tumor cohorts. Numbers indicate the significant expression differences between each pair-wise comparison. **E,** Top IPA pathways among different tumor groups, expressed as z-scores. Orange=up-regulated; blue=down-regulated. **F,** Shared gene expression subsets between and among the indicated cohorts. Red and green=number of transcripts up-regulated and down-regulated, respectively, relative to liver. **G,** Hierarchical clustering of C1 and C2A/C2B subsets ^25, 26^ of human HBs using the ten “BYN” transcripts from Table 1 that were found to be dysregulated in human tumors. **H,** Serpin E1 levels in the indicated tissues. **I,** Serpin E1 levels in plasma and cyst fluid from the indicated cohorts. Plasma and cyst fluids were diluted proportionately and 40 μg of each was subjected to SDS-PAGE. A comparable amount of lysate from one of the Δ(90)+YAP^S127A^+L30P/R34P tumors depicted in **I** was included to allow a comparison among the different samples. In order to take into account differences in the types of samples present, Ponceau acid red staining of the membrane was used to confirm protein concentrations. **J,** Gross appearance of Δ(90)+YAP^S127A^+serpin E1 tumors showing extensive necrosis (arrow). **K,** Histologic appearance of a typical section from the tumor shown in J with peri-cystic necrotic areas indicated by arrows. **L,** Immunoblots for serpin E1. Note slower mobility of exogenously expressed, epitope-tagged serpin E1.

The 821 gene expression differences between the Δ(90)+YAP^S127A^ and Δ(90)+YAP^S127A^+WT-NFE2L2 cohorts were categorized using Ingenuity Pathway Analysis (IPA). Among the four pathways with the most disparate z-scores (z>+2.8), only one (“NRF2-mediated oxidative stress response”), was considered NFE2L2-responsive (Fig. 4E and Supplementary Tables S1 and S2). In contrast, the 3584 differences between the Δ(90)+YAP^S127A^ and Δ(90)+YAP^S127A^+L30P/R34P cohorts involved 12 similarly dysregulated pathways, with three involving redox homeostasis. Finally, the 1654 differences between Δ(90)+YAP^S127A^+WT-NFE2L2 and Δ(90)+YAP^S127A^+L30P/R34P tumors showed up-regulation of the “NRF2-mediated oxidative stress response” and “glutathione redox reactions I” pathways in the latter. L30P/R34P thus dysregulated more redox target genes than WT NFE2L2.

**Table 1.**
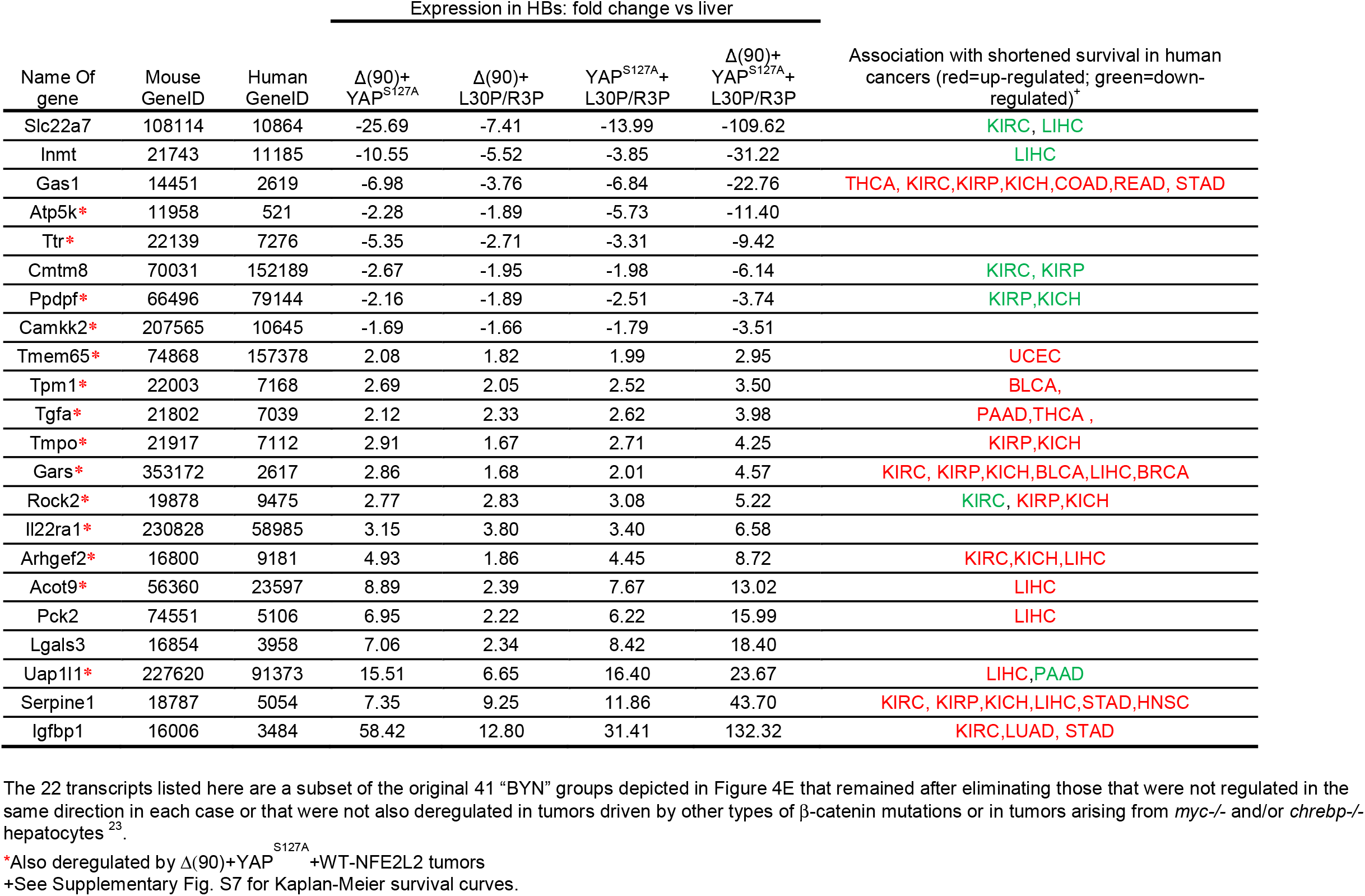
Gene responsiveness to the indicated combinations of Δ(90), YAPS127A and L30P/R34P.

ChIP-seq results from human HepG2 HB cells (https://www.encodeproject.org/data-standards/chip-seq/) indicated that 4.7-5.5% of the transcripts in Fig. 4D were orthologs of NFE2L2 direct target genes. Including data from three additional cell lines (A549, HeLa-S3 and IMR90) increased this to 13.8-22.2%. IPA profiling identified additional, variably-deregulated pathways among the tumor cohorts with the most prominent ones pertaining to Oxphos, mitochondrial dysfunction, cholesterol/bile acid synthesis and cell signaling (Fig. 4E).

41 transcripts were deregulated in all tumors with 29 always being deregulated in the same direction (Fig. 4F). 22 members of this core “BYN” subset were similarly deregulated in HBs driven by other β-catenin mutants and in slowly-growing Δ(90)+YAP^S127A^ HBs arising in *myc-/-*, *chrebp-/-* and *myc-/-+chrebp-/-* hepatocytes (Table 1)^3,24^. Δ(90)+YAP^S127A^+L30P/R34P HBs also deregulated these transcripts more than occurred with any other pair-wise combination of factors. 14 transcripts were also dysregulated in Δ(90)+YAP^S127A^+WT-NFE2L2 tumors, suggesting that they were less likely to be involved in the growth-accelerating and cystogenic properties of L30P/R34P. Together, these considerations indicated that the remaining eight coordinately de-regulated transcripts contributed to L30P/R34P’s cooperation with Δ(90) or YAP^S127A^. On average, these eight transcripts were deregulated by 47.5-fold in the Δ(90)+YAP^S127A^+L30P/R34P cohort versus 13-fold in the next highest expressing cohort (P<0.001).

Previous analyses of 24 HBs by DNA microarrays and 25 by RNAseq had identified 16 and four unrelated transcripts, respectively, whose expression correlated with survival^25,26^. In the former, tumors associated with favorable, poor and intermediate survival were designated “C1” “C2A” and “C2B”, respectively. Ten transcripts from Table 1 were differentially expressed by the C1 and C2A/C2B groups (q<0.05) with hierarchical clustering perfectly segregating the C1 and C2A/C2B groups (Fig. 4G). No transcripts in Table 1 were included in the previous panels^25,26^.

17 of the above 22 transcripts also correlated with survival in 14 cancer types from TCGA (Supplementary Fig. S7). Recurrent associations were seen with the three most common kidney cancers and HCC and some transcripts correlated with survival in multiple cancers. Of the 45 instances where expression correlated with survival, the direction of change relative to that of matched normal tissues was discordant in only one case and partially discordant in two others (Table 1).

The Cancer Hallmarks Analytics Tool (http://chat.lionproject.net/) showed associations of the above 22 transcripts with known cancer-promoting/enabling features^33,34^. 16 transcripts were associated with one or more hallmarks and eight were associated with five-nine (Table 2). By comparison, the associations of five well-known oncogenes and tumor suppressors, with broad human cancer associatrions, ranged from five-eight. BYN transcript levels therefore frequently correlate with the properties, behaviors and lethality of multiple cancers, including HBs.

**Table 2.**
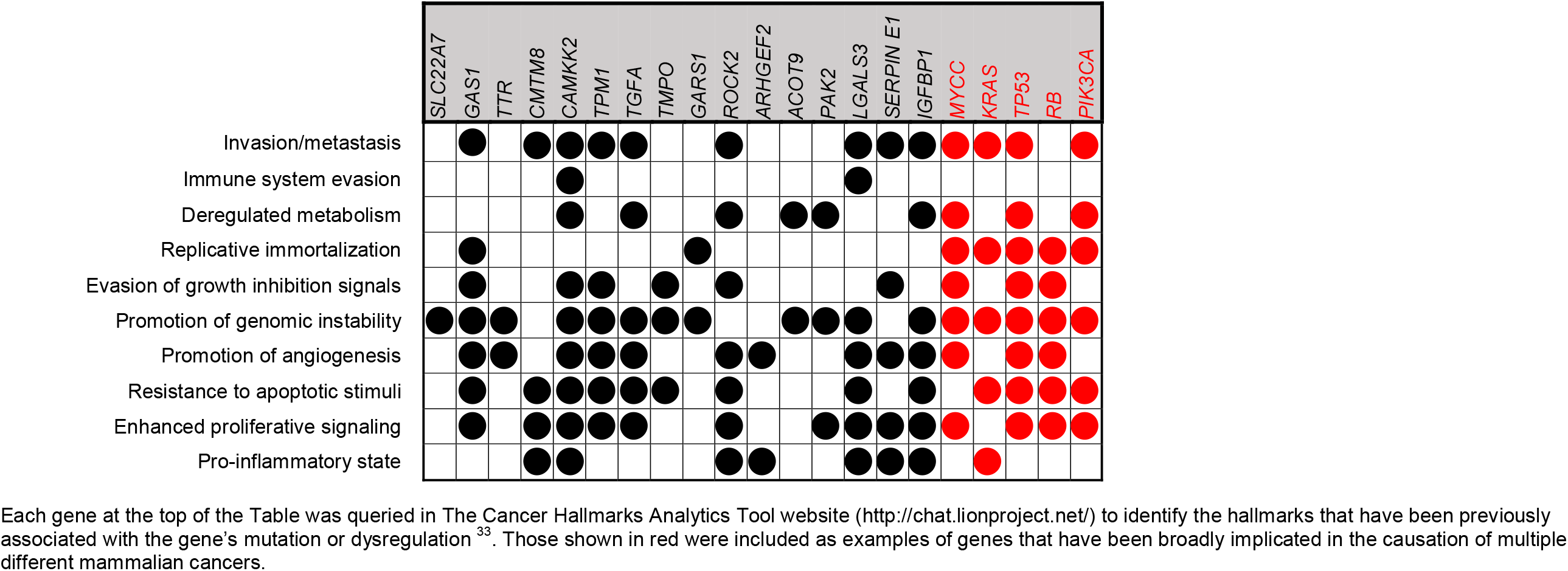
Association of relevant “BYN” transcripts with The Hallmarks of Cancer.

Using data from HepG2 HB cells and eight additional human cell lines in the ENCODE v.5 CHIP-seq data base, the promoters of 20 of the above 22 BYN genes were found to contain at least one *bona fide* binding site for β-catenin or YAP/TAZ although only two genes contained documented sites for NFE2L2 (Supplementary Figs. S8 and S9). Thus, many of the BYN genes are either indirect NFE2L2 targets or harbor biding sites elsewhere.

The *serpine1* gene promoter contained the largest number of binding sites for all three transcription factors (Supplementary Fig. S9B). Serpin E1 levels were particularly high in Δ(90)+YAP^S127A^+L30P/R34P tumors (Fig. 4H) and agreed with RNAseq results (Table 1). Cyst fluid and plasma from tumor-bearing mice also contained serpin E1 whereas plasma from mice bearing Δ(90)+YAP^S127A^ tumors contained low-undetectable levels (Fig. 4I).

Serpin E1 co-expression affected neither the survival of Δ(90)+YAP^S127A^ tumor-bearing mice nor promoted cysts (Fig. 1A). However, the tumors were necrotic and expressed serpin E1 (Fig. 4J-L). Serpin E1 deregulation thus recapitulated the extensive necrosis associated with L30P/R34P over-expression but not the accelerated growth or cystogenesis.

### *NFE2L2* mutations, CNVs and over-expression in HB and other cancers

Our RNAseq analyses of 45 murine tumors generated by nine β-catenin variants identified only L30P, which was previously described in human HBs^3^. We therefore queried RNAseq or exome-seq data from 194 previously reported primary HBs and cell lines (https://cancer.sanger.ac.uk/cosmic) ^7, 26, 35^ and identified nine NFE2L2 missense mutations (frequency=3.5%) (Supplementary Table S3).

In contrast to missense mutations, up to 50% of HBs harbor NFE2L2 CNVs^35^. We quantified *NFE2L2* gene copy numbers in 22 primary HBs from our own institution and identified three (13.6%) with 4.0-6.4-fold increases in copy number (Supplementary Fig. S10A). Up to 25% of other human cancers also showed *NFE2L2* gene amplification (https://www.mycancergenome.org/content/gene/nfe2l2/ and Supplementary Fig. S10B). However, an association between low levels of NFE2L2 and KEAP1 transcripts and favorable survival was seen only in HCCs (Supplementary Fig. S10C and D).

Transcript and protein levels often correlate poorly, particularly in cancer^36^. This is especially true for NFE2L2 whose activation occurs post-translationally^10,11,27^. Evaluating survival based on transcript levels (Supplementary Fig. S10C and D) might therefore not fully capture the magnitude of NFE2L2’s influence. A more meaningful way to identify NFE2L2 deregulation that reflects the protein’s redox-dependent function might be to survey tumors for NFE2L2 target gene expression. We did this for 46 direct targets from the Ingenuity Pathway Analysis Knowledge Base (Supplementary Table S4) in three cancer types from TCGA which preliminary analysis indicated as having high incidences of *NFE2L2* CNV or mutation. NFE2L2 transcript levels in these tumors were similar to those in matched normal tissues and did not correlate with target gene transcript levels. As a group however, these transcripts were significantly up-regulated (1.36-1.63-fold) (Fig. 5A-C) and the survival of individuals with tumors expressing the highest levels was significantly shorter (Figs. 3A and 5D-F). This suggests that NFE2L2 target gene products suppress ROS and other electrophiles that compromise tumor aggressiveness^8,19,35^.

**Figure 5.**
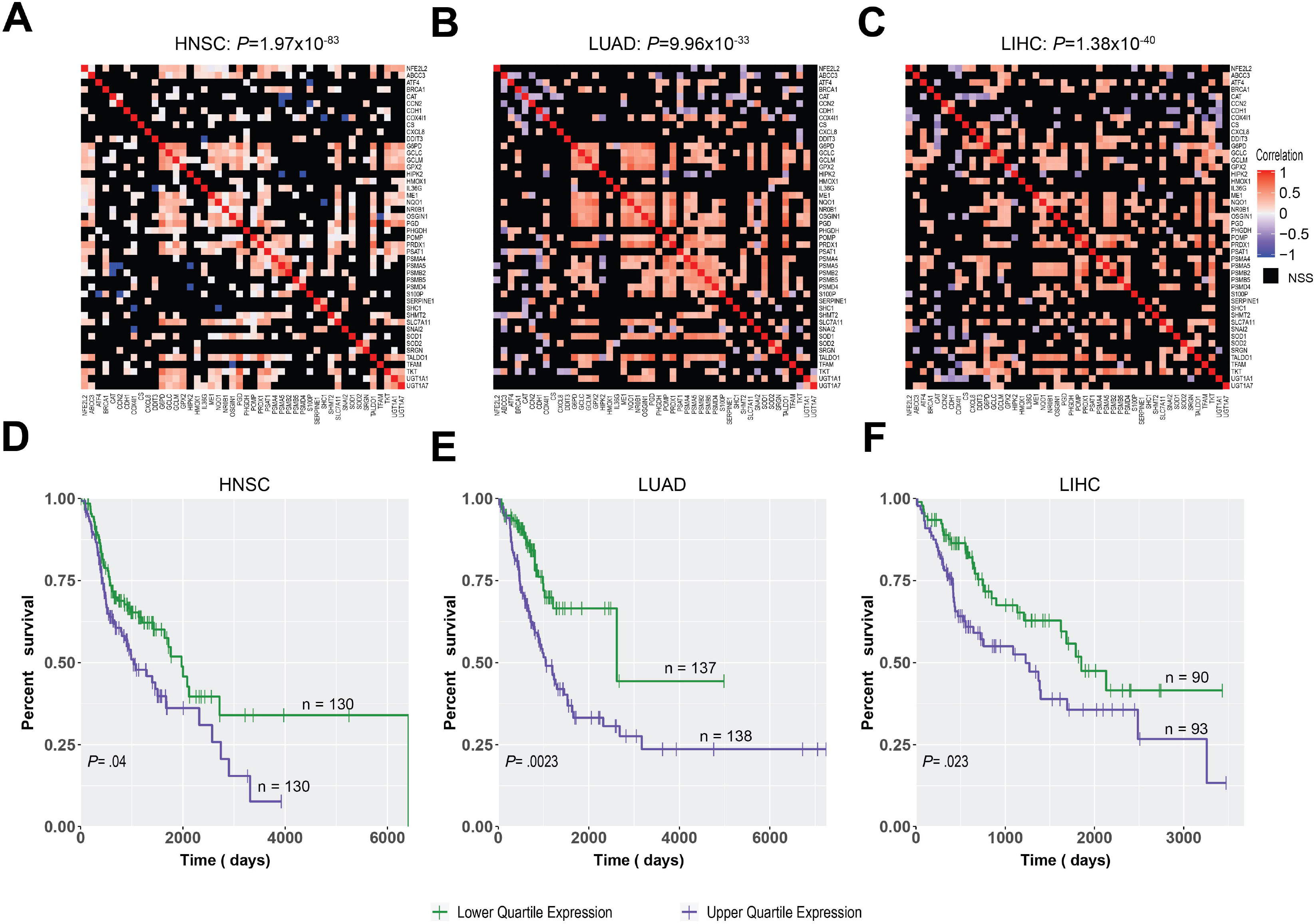
Correlation between NFE2L2 and its target genes in three human cancer types. **A-C,** Correlation matrices of NFE2L2 and 46 NFE2L2 target gene transcripts (Supplementary Table S4) from three of the tumor groups depicted in Supplementary Fig. S10B (**A**=head & neck squamous cell cancer [HNSC], **B**=lung adenocarcinoma [LUAD]), **C**=HCC [LIHC]. The preponderance of positive correlations is apparent and was assessed using a binomial test P values shown at the top of each panel). **D-F,** Long-term survival of patients whose tumors are profiled in **A-C**. Expression levels of the 46 NFE2L2 target genes (Supplementary Table S4) were averaged across all samples for each cancer type. Survival differences between the two quartiles of individuals whose tumors expressed the highest and lowest levels of these transcripts were determined using Kaplan-Meier survival and were assessed using log-rank tests.

## Discussion

L30P and R34P, but not WT-NFE2L2, markedly accelerate murine liver tumorigenesis, supporting the idea that NFE2L2-KEAP1 dysregulation suppresses the proliferative limitations imposed by oxidative, electrophilic and metabolic stresses^10–13^. Depending on context and timing, this may have opposite outcomes^9^. For example, it can reduce ROS-mediated oncogenic lesions that initiate tumorigenesis. At later times, it can increase tolerance to oncoproteins and facilitate tumor evolution, expansion and therapy resistance. Contributing non-canonical functions of NFE2L2 targets might include the regulation of apoptosis, metabolism, angiogenesis and chemotherapeutic drug detoxification^19,20,37^. Support for this model, however, has derived largely from *in vitro* studies or from molecularly heterogeneous tumor xenografts^37–39^. Our findings show that L30P/R34P not only accelerate Δ(90)+YAP^S127A^-mediated tumorigenesis but are directly transforming when co-expressed with either oncoprotein, thus indicating that some HBs could be NFE2L2-driven when only one arm of the β-catenin/Hippo axis is de-regulated^2,4^. Further examination of other cancers harboring *NFE2L2* mutations or CNVs may reveal unrecognized co-dependencies with other oncoproteins or tumor suppressors.

A unique feature of Δ(90)+YAP^S127A^+L30P/R34P or R582W+YAP^S127A^+L30P/R34P HBs is widespread cystogenesis (Fig. 1B-E,G and Supplementary Figs. S2-4). Tumor vasculature is sometimes comprised of actual tumor cells or tumor cell-derived endothelium^40,41^. However, the numerous, large and bloodless cysts we observed do not appear to be of vascular origin. “Peliosis hepatis” is a rare condition associated with epithelial- or endothelial-lined cyst-like lesions but these are typically blood-filled and less abundant. The cysts we observed are more reminiscent of those occasionally seen in human HBs^21^, were independent of tumor growth rates and arose only in the context of β-catenin+YAP^S127A^+L30P/R34P co-expression.

In contrast to its cytoplasmic location in cultured cells, nuclear WT-NFE2L2 (Fig. 2B) may reflect the oxidized cytoplasm of tumor cells and/or the hepatocyte’s abundance of dietary electrophiles^34,42^. These chronic stresses could maintain high levels of oxidized KEAP1 with any reduced residuum being responsible for lowering WT-NFE2L2 levels (Fig. 2A). The 821 transcript differences between Δ(90)+YAP^S127A^ tumors and Δ(90) +YAP^S127A^+WT-NFE2L2 tumors (Fig. 4D and Supplementary Table S2) is testimony to the consequences of even small perturbations in the NFE2L2-KEAP1 balance. This likely explains why either mutations or CNVs increase the fractional nuclear concentration of NFE2L2 with similar consequences^8,38^.

Δ(90)+YAP^S127A^+L30P/R34P tumor metabolism resembled that of Δ(90)+YAP^S127A^ HBs including lower Oxphos and mitochondrial mass and higher pyruvate consumption compared to livers (Figs. 2C-I and 3C-I)^3,23^. In contrast, they variably increased pPDHα1, Glut1, Glut2, PKM2 and PFK-L (Fig. 3F and J). Higher activity of PDH, which links glycolysis and Oxphos, may allow tumors to compensate for lower FAO rates by maximizing AcCoA synthesis in the face of ongoing Warburg respiration^3,23^. This dynamic behavior underscores the balancing of FAO and glycolysis via the Randle cycle^43^, which allows for new membranes to be synthesized from pre-existing fatty acids and minimizing the need for *de novo* synthesis from more primitive precursors.

Despite alterations of glutaminolysis pathway enzymes (Fig. 3J), Δ(90)+YAP^S127A^+L30P/R34P tumors also decreased glutamine-driven Oxphos, which is used by some tumors to maintain α-ketoglutarate pools^43^. Collectively, these studies indicate that metabolic demands of these tumors were addressed by reprogramming glycolysis and the TCA cycle.

Δ(90)+L30P/R34P HBs were unique in their down-regulation of Glut1 and PKM2 and their up-regulation of pPDHα1, Glut4 and PFK-L, a rate-limiting glycolytic enzyme^44^. This complex relationship suggested a potentially higher rate of glycolytic flux into mitochondria while simultaneously reducing the accumulation of anabolic precursors as a result of slower tumor growth rates. However, the mitochondrial mass and Oxphos of these tumors exceeded even that of normal livers (Fig. 3C-E,I and Supplementary Fig. S6). Together, these findings underscored a reduced reliance on Warburg respiration and a return to a more liver-like metabolic profile.

In further contrast, the metabolic behavior of YAP^S127A^+L30P/R34P tumors more closely resembled that of Δ(90)+YAP^S127A^ tumors (Fig. 3C-H) although with different expression patterns of the above proteins. Notable differences included the virtual absence of PC, the consequence of which might be the enhanced catalysis of pyruvate to AcCoA.

Tumors with pair-wise Δ(90), YAP^S127A^ and L30P/R34P combinations allowed for the assignment of distinct, albeit general, functions for each factor and the identification of key individual transcriptional repertoires. Thus, Δ(90), but not YAP^S127A^, appears to promote increased mitochondrial mass and a more metabolically active state in response to all tested substrates. Yet, the metabolic behaviors of Δ(90)+YAP^S127A^ and YAP^S127A^+L30P/R34P tumors (Fig. 3C-H) suggest that YAP^S127A^ was dominant over Δ(90), which tended to suppress these more exaggerated responses.

Unbiased RNAseq permitted the identification of specific transcripts associated with each 2-3 member combination of Δ(90)/YAP^S127A^/L30P/R34P (Fig. 4B-F). Importantly, it revealed a common 22 transcript subset deemed most likely to explain the accelerated growth rates, cystogenesis and necrosis of Δ(90)+YAP^S127A^+L30P/R34P tumors (Table 1). Further evaluation provided additional evidence for its importance to HBs and other cancers. First, all 22 transcripts were altered in the same direction in each HB cohort (Table 1 and Fig. 4F). Second, they were expressed in tumors with disparate growth rates generated by different β-catenin mutants^3,23^. Third, they were more deregulated in Δ(90)+YAP^S127A^+L30P/R34P tumors (Table 1). Fourth, ten of the transcripts perfectly identified human HB groups with different prognoses (Fig. 4G)^25,26^. Lastly, 16 of the transcripts were associated with cancer-promoting or enabling functions (Table 2).

Among the most intriguing of the 22 transcripts was that encoding serpin E1/plasminogen activator inhibitor-1, a serine protease inhibitor with roles in tumor growth, metastasis, angiogenesis and matrix remodeling in addition to canonical functions in fibrinolysis^45^. Serpin E1 transcripts were the second most highly up-regulated in β-catenin+YAP^S127A^+L30P/R34P tumors (Table 1) and the *serpine1* promoter was unique in containing multiple *bona fide* response elements for each transcription factor (Supplementary Figs. S8 and S9). We also extended previous observations that serpin E1 levels correlate with unfavorable survival in human cancers (Supplementary Fig. S7)^46,47^. High levels of circulating serpin E1 are associated with polycystic ovary syndrome and serpin E1’s enforced expression in murine ovaries is cystogenic^48,49^. Serpin E1 co-expression in β-catenin+YAP^S127A^ HBs did not accelerate growth rates or promote cystogenesis but did recapitulate the extensive necrosis of β-catenin+YAP^S127A^+L30P/R34P HBs (Figs. 1D and 4J and K). Other transcripts therefore likely impact tumor growth rates and cystogenesis.

Our findings demonstrate that NFE2L2 mutants alter redox balance in β-catenin+YAP^S127A^ HBs and increase growth, cystogenesis and necrosis. The unanticipated oncogenicity of L30P/R34P when co-expressed with β-catenin or YAP^S127A^ also demonstrated their direct role in transformation *in vivo* and unequivocally establish NFE2L2 as an oncoprotein that can be activated by mutation, over-expression or other factors that perturb NFE2L2:KEAP1 balance. This work also identified key transcripts that likely mediate these unique features and directly contribute to tumorigenesis. The high rate of NFE2L2-KEAP1 axis dysregulation in cancer and its prognostic implications can now be better appreciated in light of these findings.

## Supporting information

Supplementary Methods, Figures S1 to S10, Tables S1 to S5 and References

