## Supplementary Methods, Figures S1 to S10, Tables S1 to S5 and References for "Patient-Derived Mutant Forms of NFE2L2/NRF2 Drive Aggressive Murine Hepatoblastomas"

[1]Division of Hematology/Oncology, UPMC Children's Hospital of Pittsburgh; [2]Tsinguhua University School of Medicine, Beijing, People's Republic of China; [3]The University of Pittsburgh School of Medicine; [1,4]Central South University Xiangya University Medical School, Changsha, People's Republic of China; [5]Department of Pathology, Cincinnati Children's Hospital, Cincinnati, OH; [6]The Department of Microbiology and Molecular Genetics, UPMC; [7]The Hillman Cancer Center, UPMC; [8]The Pittsburgh Liver Research Center, Pittsburgh, PA.

Corresponding author:
Edward V. Prochownik, M.D., Ph.D.


**This PDF file includes:**

Supplementary Methods
Figures S1 to S10
Tables S1 to S5
References

#### Supplementary Methods

**Oxphos measurements.** OCRs were determined on partially purified mitochondrial suspensions in MiR05 buffer following the addition of cytochrome c (10 μM), malate (2 mM), pyruvate (5 mM), ADP (5 mM) and glutamate (10 mM) to initiate electron transport chain activity via Complex I (3,25). Succinate was added to a final concentration of 10 mM to allow the sum of Complexes I + II activities to be measured. Rotenone (0.5 μM final concentration) was added to inhibit Complex I and to allow the individual contributions of Complex I + Complex II to be confirmed. All activities were normalized to total protein. To measure β-FAO, palmitoyl-CoA and L-carnitine (3 μM and 10 μM, respectively) were added to reactions already primed with ADP and malate.

**Tumor studies.** Animal studies were performed in compliance with the Public Health Service Policy on Humane Care and Use of Laboratory Animals Institute for Laboratory Animal Research Guide for Care and Use of Laboratory Animals. They were also approved by the Institutional Animal Care and Use Committee at the University of Pittsburgh.

**Oxphos measurements.** OCRs were determined on partially purified mitochondrial suspensions in MiR05 buffer following the addition of cytochrome c (10 μM), malate (2 mM), pyruvate (5 mM), ADP (5 mM) and glutamate (10 mM) to initiate electron transport chain activity via Complex I (1-3). Succinate was added to a final concentration of 10 mM to allow the sum of Complexes I + II activities to be measured. Rotenone (0.5 μM final concentration) was added to inhibit Complex I and to allow the individual contributions of Complex I + Complex II to be confirmed. All activities were normalized to total protein. To measure β-FAO, palmitoyl-CoA and L-carnitine (3 μM and 10 μM, respectively) were added to reactions already primed with ADP, malate and succinate.

**Oxidative stress.** Tumors expressing cyto-roGFP or mito-roGFP (4) were minced in D-MEM+10% FBS and plated in 100 mm plastic tissue culture dishes. Cells were allowed to attach to 100 cm tissue culture plate and were maintained until achieving ~80% confluence. Cells were then trypsinized and re-seeded into 35 mm glass bottom plates (MatTek, Corp., Ashland, MA), allowed to attach for two-three days and visualized in real time in a temperature- and $CO_2$-controlled environment using a LSM710 laser scanning confocal microscope (Carl Zeiss, Munich, Germany). Emission signal intensity ratios (488/405 nm) from regions of comparable brightness was used to determine baseline redox states of each compartment.

**TaqMan Assays.** Final PCR primer concentrations were 250 nM each and each TaqMan probe concentration was 300 nM. NFE2L2 primer sequences were: (Fwd): 5'-TCATCTACAAACGGGAATGTCT-3', (Rev): 5'-GTTGCCCACATTCCCAAATC-3', TaqMan probe: 5'-56-FAM/AGATGCTTT/ZEN/GTACTTTGAT-3'. Two nuclear genes were used for controls, namely glyceraldehyde-3-phosphate dehydrogenase (GAPDH) and Ribonuclease P RNA Component H1 (RPPH1). GAPDH primer sequences were: (Fwd): 5'-CCTAGGGCTCACATATTC-3', (Rev): 5'-CGCCCAATACGACCAAATCTA-3', TaqMan probe: 5'-/5Cy5/TCCTCATGC/TAO/CTTCTTGCCTCTTGT /3IABRQSP/3'. RPPH1 PCR primer sequences were: (Fwd): 5'-TCTGGCCCTAGTCTCAGACCTT-'3 , (Rev): 5'-GAGCTGAGTGCGTCCTGTC-3', TaqMan probe: 5'-56-FAM/CCAAGGGAC/ZEN/ATGGGAGTG-3'. Reactions contained 50 ng of total DNA and were performed on a CFX96 Touch$^{TM}$ real-time PCR detection system (Bio-Rad, Inc., Hercules, CA) Amplification conditions were 95 °C for 10 s followed by 40 cycles at 95 °C for 15 s. and 60 °C for 60 s.

Mitochondrial DNA (mtDNA) content was quantified using a TaqMan-based approach that amplified a 90 bp segment of the D-loop region as previously described (1,3). The results were normalized to a control TaqMan reaction that amplified a 73 bp region of the nuclear apolipoprotein B gene. Reactions contained 10 ng of total DNA and were performed on a CFX96 Touch™ real-time PCR detection system (Bio-Rad, Inc., Hercules, CA) using the conditions: 95 °C for 10 s, 40 cycles at 95 °C for 15 s. followed by 60 °C for 60 s.

**Immunoblotting.** Tissue lysates were prepared in SDS-lysis buffer as previously described (1,3). Immunoblots were developed using SuperSignal™ West Pico Chemiluminescent Substrate kit (Thermo Fisher, Waltham, MA). Antibodies, the vendors from which they were obtained and the conditions used are listed in Supplementary Table S5.

For subcellular fractionation nuclear and cytoplasmic fractionations were performed on ~100 mg tissue fragments using a Subcellular Protein Fractionation Kit as previously described (3) according to the supplier's directions (Thermo-Fisher).

**RNAseq.** RNA purification was performed using Qiagen RNAeasy columns according to the directions of the supplier. An Agilent 2100 Bioanalyzer (Agilent Technologies, Foster City, CA) was used to evaluate integrity and only samples with RIN values >8.5 were used. Samples were prepared for paired-end sequencing using an NEB NEBNext Ultra Directional RNA Library Prep kit (New England Biolab, Beverly, MA) and sequencing was performed on a NovaSeq 600 Instrument (Illumina, Inc., San Diego, CA) by Novagene, Inc. (Sacramento, CA). Raw and processed original data are accessible through GEO (accession number: GSE157623) (https://www.ncbi.nlm.nih.gov/geo/query/acc.cgi?acc= GSE157623). Differentially-expressed transcripts were identified using DeSEq2, CLC Genomic Workbench v. 12.0 (Qiagen), and EdgeR and were analyzed using the Galaxy platform (https://galaxyproject.org/use/). For DeSeq2 and EdgeR, FASTQ file reads were mapped against the GRCm38.p6 mouse reference genome using STAR (https://github.com/alexdobin/STAR/releases) version 2.5.2. The output files were analyzed by featureCounts (http://bioinf.wehi.edu.au/featureCounts/) to quantify transcript abundance. Only transcripts whose differential was demonstrated by all three methods are reported and only after significance was adjusted for false discovery using the Bonferonni–Hochberg correction ($q<0.05$). Ingenuity Pathway Analysis (IPA, Qiagen) was used for pathway identification. The following parameters were recorded for each pathway: *p* value, ratios of dysregulated transcripts to all transcripts associated with that pathway (ratio), and predicted pathway activation, inhibition and indeterminate (*Z*-score).

**Target gene promoter analysis.** Human and mouse genomes were accessed through the BSgenome.Hsapiens.UCSC.hg38 and BSgenome.Mmusculus.UCSC.mm10 R packages, respectively. Gene position information was accessed for human and mouse respectively using the TxDb.Hsapiens.UCSC.hg38.knownGene and TxDb.Mmusculus.UCSC.mm10.knownGene packages respectively. The flank function was used to query the upstream 5000 bp promoter regions for each gene and the searchSeq function of the TFBSTools R package was used to assess the binding sites in these flanking regions using the aforementioned position frequency matrices. Binding position frequency matrices were queried in R from the JASPAR database using the package JASPAR2018 and converted to position weight matrices using the toPWM function of the TFBSTools R package. Sites with a binding scores >90% were accepted as putative binding sites.

Chip-Seq Data were accessed from the ENCODE v.5 database (https://www.encodeproject.org/) for the terminal transcription factors (TF) of the NFE2L2 (ARE sites), Wnt/β-Catenin (Tcf/Lcf sites), and Hippo/Yap (TEAD sites) pathways. Data were only included from cell lines with readily available IDR thresholded peaks identified, and p-values derived from more than one experimental replicate. The cell lines from which data was obtained included HepG2, A549, Hela-S3, IMR90, GM12867, HCT-116, HEK293T, K562 and PANC-1. The RTrackLayer R package was used to process the data and the R packages BSgenome.Hsapiens.UCSC.hg38 and TxDb.Hsapiens.UCSC.hg38.knownGene were used to align the Chip-Seq data to the promoters of the genes of interest. For figures summarizing the binding of more than one TF, the p-value plotted was the maximum of any of the TFs at that base-pair.

**Dysregulation of NFE2L2 and its direct target genes in human cancers.** RNAseq data from 141 human HB samples (5,6) were searched for NFE2L2 mutations using CLC Genomics Workbench 11. The data were uploaded and a track from reference CDS annotation of gene *NFE2L2* was generated from the “Homo_sapiens_refseq_GRCh38.p11_o_Genes” track. Amino acid changes within each sample were found using Basic variant detection: “Toolbox>Resequencing Analysis>Functional Consequences>Variant detection>Basic variant detection” followed by “Amino Acid Changes”: “Toolbox>Resequencing Analysis>Functional Consequences>Amino Acid Changes”. NFE2L2 mutations from 116 human HBs from the COSMIC Database of were identified by performing a search using “Tissue Distribution>Liver>Sub-Histology> Hepatoblastoma”._*NFE2L2* gene CNVs across multiple cancers were identified by analyzing TCGA genomic data in Genomic Data Commons (GDC).

NFE2L2 and KEAP1 transcript expression data were downloaded from the TCGA PANCAN dataset as FPKM, converted to TPM, and filtered to contain only data from HCCs. Samples were defined as having high (top 40%) or low (bottom 40%) levels of KEAP1 and NFE2L2, thus allowing data to be independently categorized according to the expression of each tumor’s transcripts. Survival analysis for each group was performed using the Survival package in R, displayed as Kaplan-Meier curves and compared pairwise for log-rank p values. This procedure was repeated for OV, LUSC, ESCA, HNSC, SARC and LUAD tumors.

To determine the relationship between NFE2L2 expression and its direct targets, we used IPA to identify 46 promoters that contained established NFE2L2-binding ARE elements. Expression data (TPM) for these genes and NFE2L2 itself were accessed from the GDC TCGA dataset available on the UCSC Xenabrowser (xena.ucsc.edu) from cohorts with the highest levels of NFE2L2 CNVs. Natural log-fold up- or down-regulation for each transcript was computed relative to that in matched normal tissues from the same individual. For visualization purposes, the pairwise correlation matrices of differential values were plotted as a heat map for each cohort, first with transcripts in an arbitrary order that was the same across cohorts, and then as hierarchically clustered heat maps. In all cases only significant correlations were expressed in color with any remaining ones being blacked out. The default settings of the R “ComplexHeatmap” package were used for hierarchical clustering. The predominance of positive correlations was assessed using a binomial test.

**Statistical analyses.** Survival data for patients in TCGA, were analyzed with GraphPad Prism 7 (GraphPad Software, Inc., San Diego, CA). Depending on the tumor type and “BYN” member transcript being analyzed (Fig. 4E), different expression level cut-offs were used to define the intragroup subsets with the highest and lowest levels of expression (e.g., higest vs. lowest 50%, quartile, etc.). Kaplan-Meier survival curves were generated by Log-rank (Mantel-Cox) test. One-way ANOVA was applied for multiple comparisons using Fisher’s least significant difference test. Student’s 2-tailed t test was used for comparing differences between 2 groups.

#### Supplementary Figures

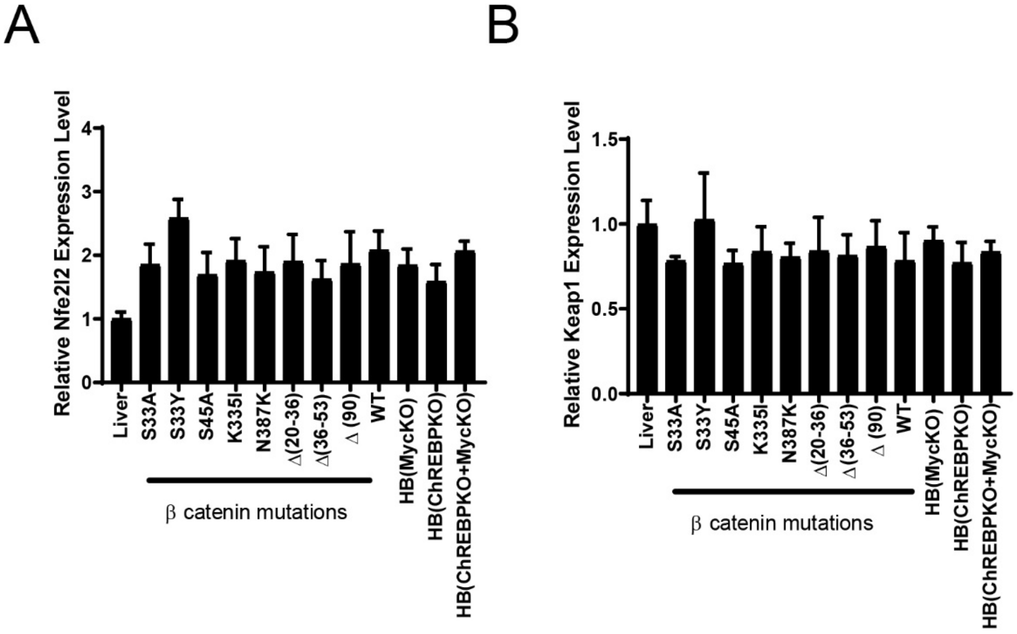


**Supplementary Figure S1.**
NFE2L2 and KEAP1 transcript levels in murine HBs. **A,** NFE2L2 transcript levels were quantified from previously reported RNA-seq data obtained from murine HBs generated by the enforced hepatic overexpression of $YAP^{S127A}$ and eight missense or in-frame deletion mutants of β-catenin or WT β-catenin (3). Transcripts were also quantified in Δ(90) β-catenin-generated HBs arising in myc-/-, chrebp-/- and myc-/- x chrebp-/- hepatocyte backgrounds (25,56). All transcript levels are expressed relative to those in normal liver (n=5 samples/group). **B,** KEAP1 transcript levels in the tissues shown in **A.**

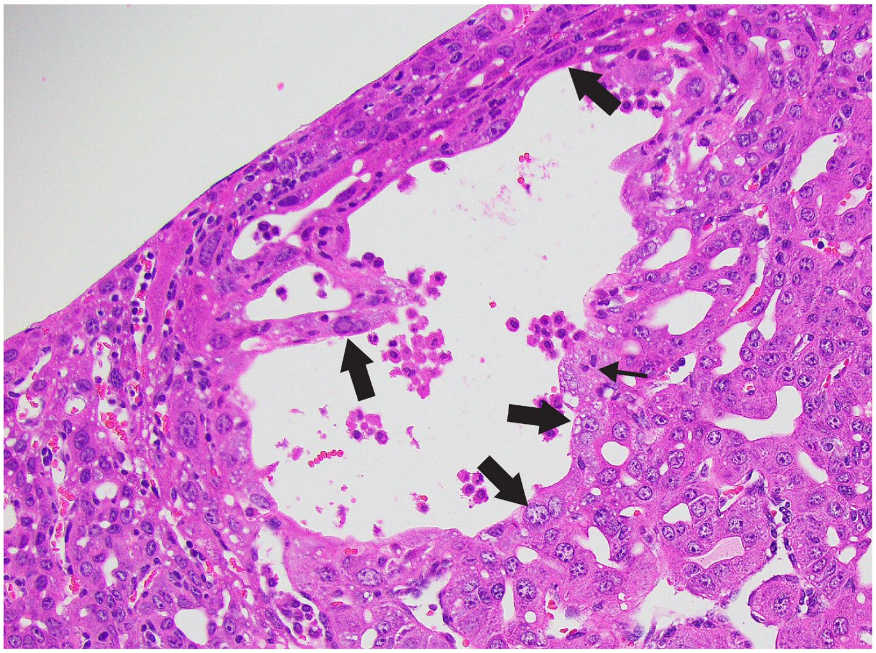

**Supplementary Figure S2.**
Cyst structure. High power magnification of an H&E-stained section showing the lumen of a typical cyst in an HB generated by the combination of $\Delta$(90)+YAP$^{S127A}$+L30P (Fig. 1**D**). The cyst is lined with cells indistinguishable from those lying deeper in the parenchyma and includes some with the multiple nuclei typical of hepatocytes (thick arrows). A presumptive mitotic tumor cell is indicated by the thin arrow.

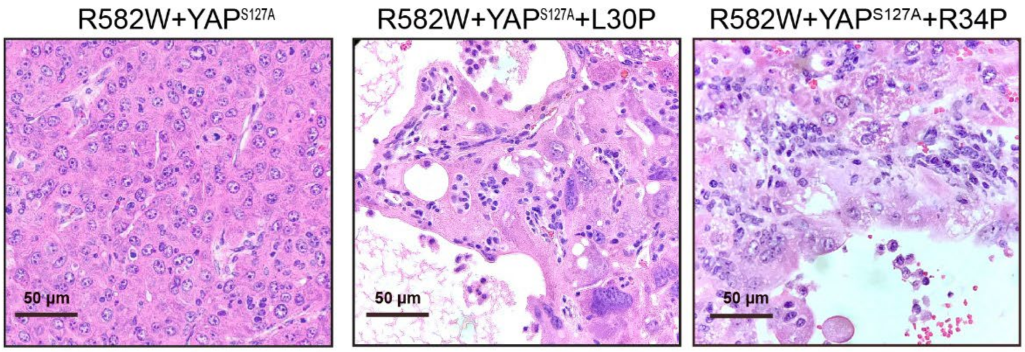


**Supplementary Figure S3.**
H&E-stained sections of tumors generated by R582W+YAP$^{S127}$+/-L30P/R34P. Note that all tumors more closely resemble HCCs than HBs but that those expressing L30P or R34P show more cellular atypia, a greater number of mitoses and cysts. See Zhang et al (3) for additional images of R582W+YAP$^{S127}$ tumors.

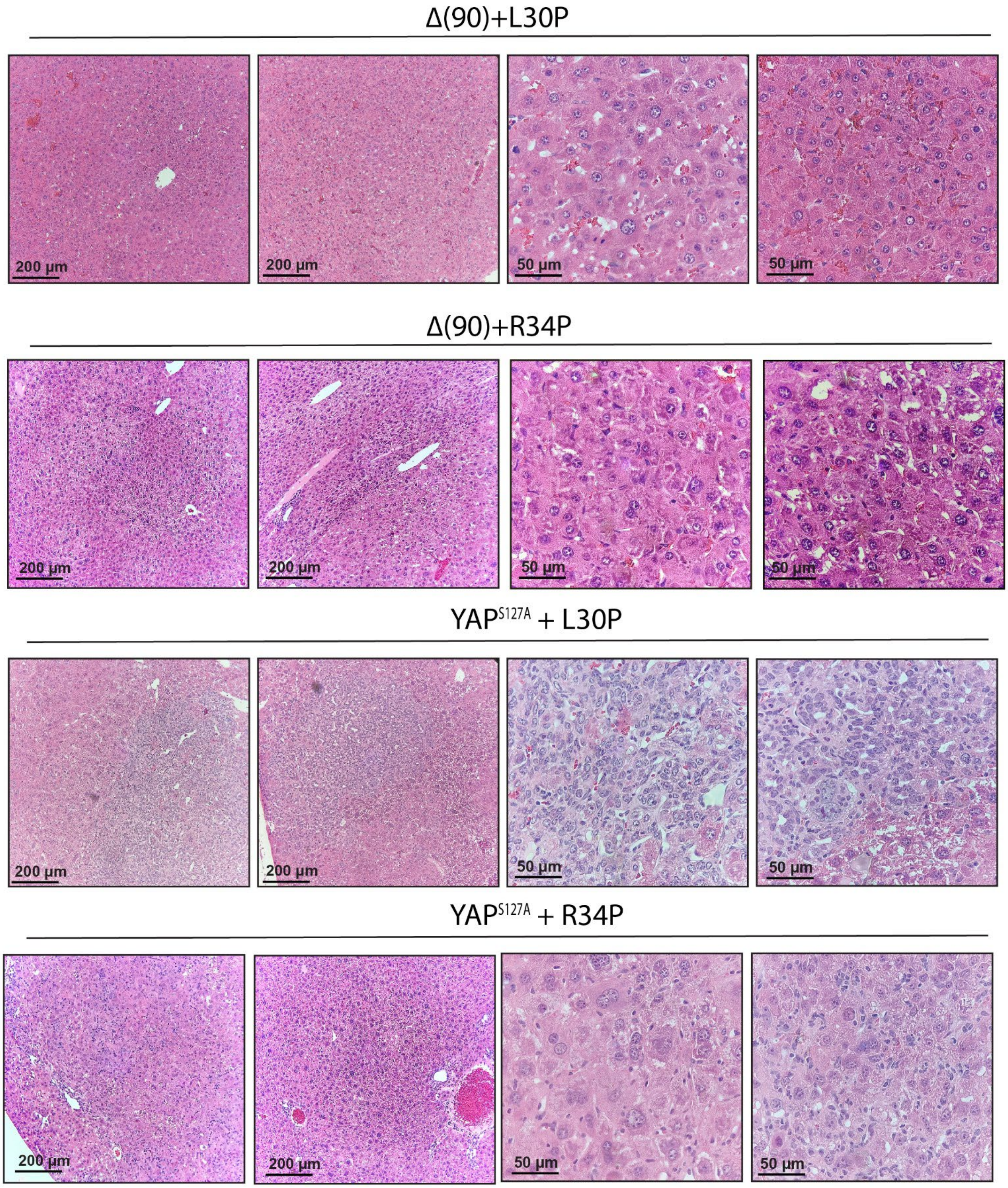


**Supplementary Figure S4.**
Additional H&E images of representative tumors.

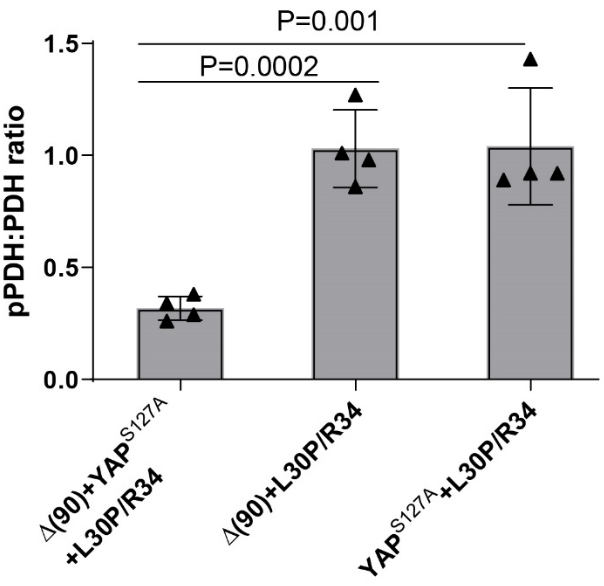


**Supplementary Figure S5.** Quantification of PDH and pPDH immunoblot results from Fig. 3**J**.

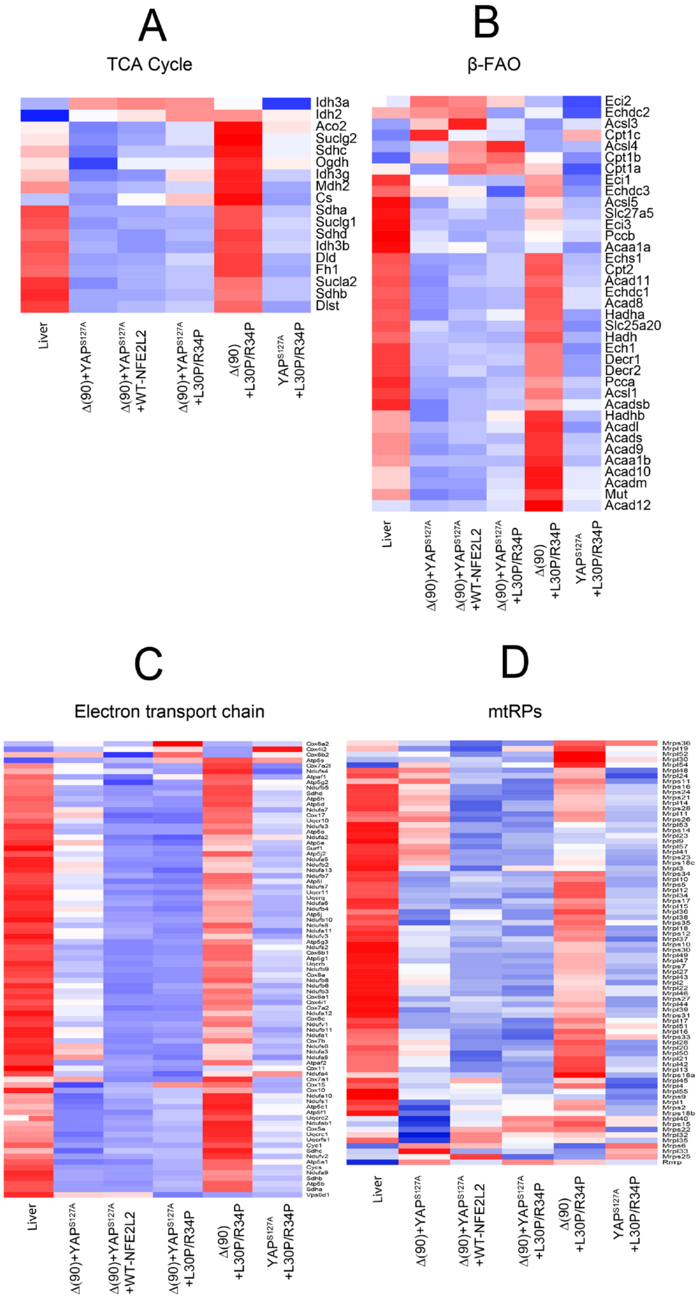


**Supplementary Figure S6.**
Differential expression of transcripts involved in mitochondrial function. Control livers or tumors from each of the indicated cohorts (n=5/group) were subjected to RNAseq and analyzed for expression of transcripts encoding proteins in the following pathways or structures. **A,** TCA Cycle, **B,** β-FAO. **C,** Electron transport chain, **D.** mtRPs

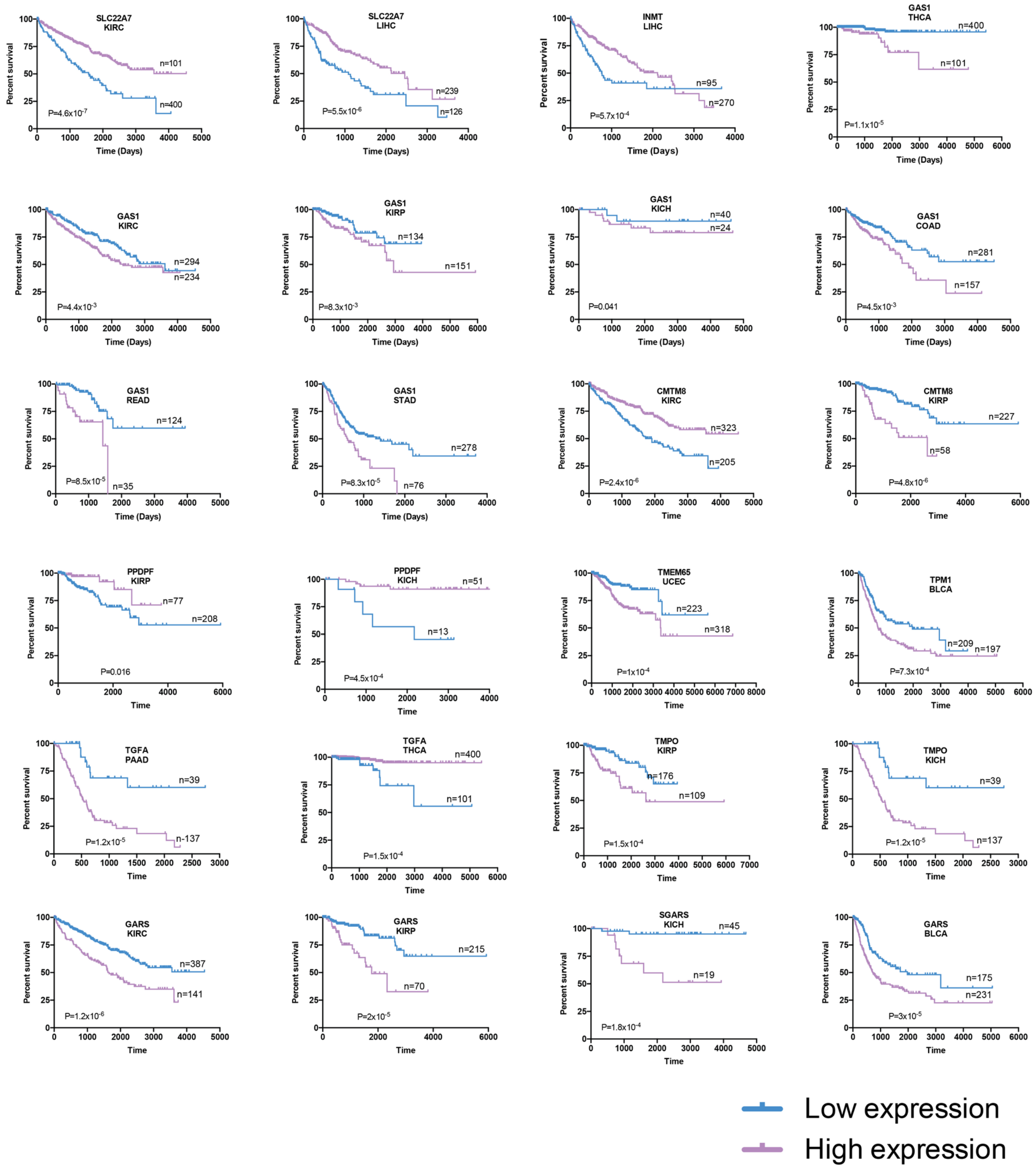


**Supplementary Figure S7**

Correlation between expression levels of transcripts listed in Table 1 and survival in select human cancers. Each depicted tumor type was divided into two groups displaying the highest and lowest expression of the indicated transcript. Standard Kaplan-Meir survival curves for each group were then generated and P values were determined by a standard rank test.

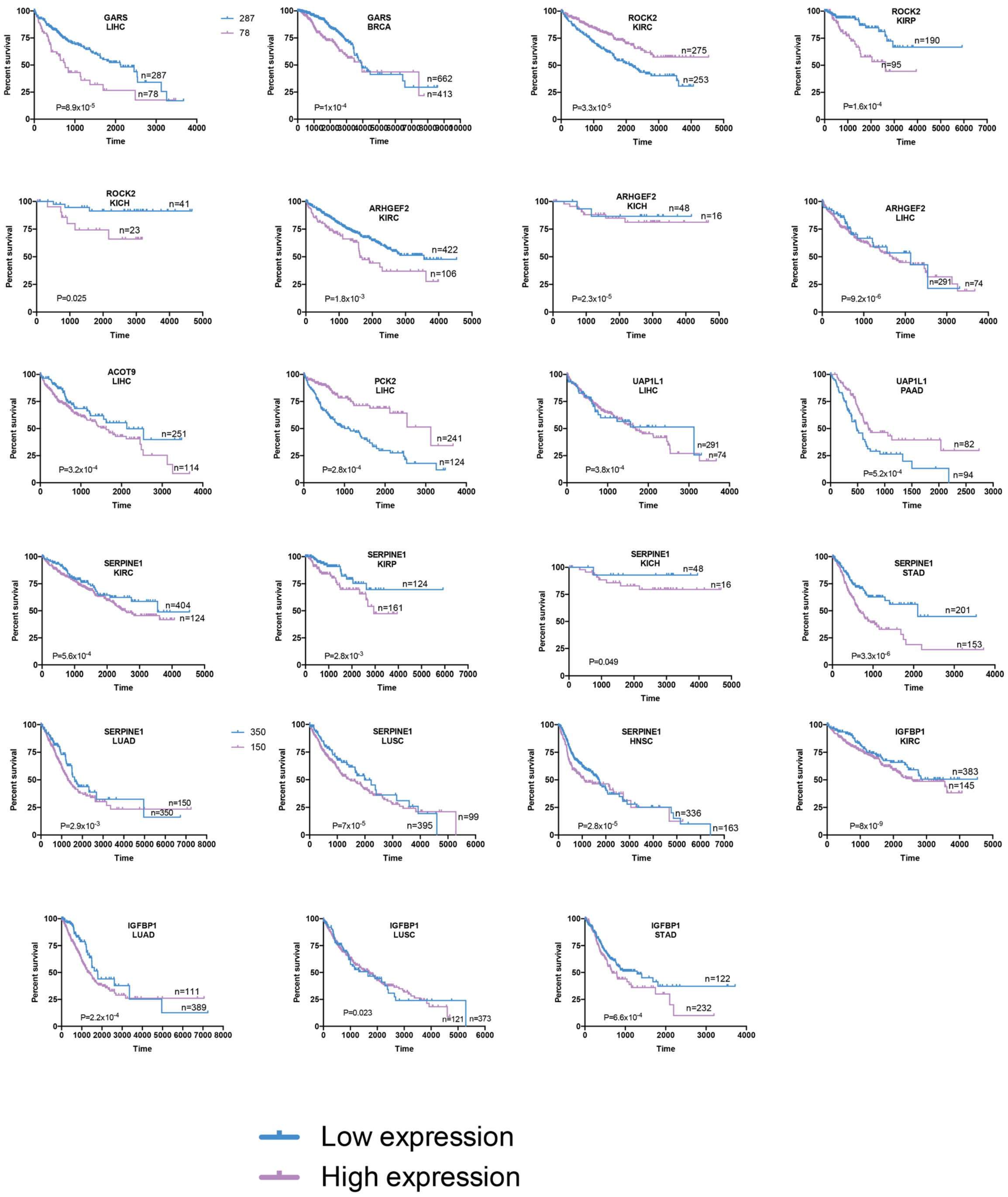


**Supplementary Figure S7 (continued)**

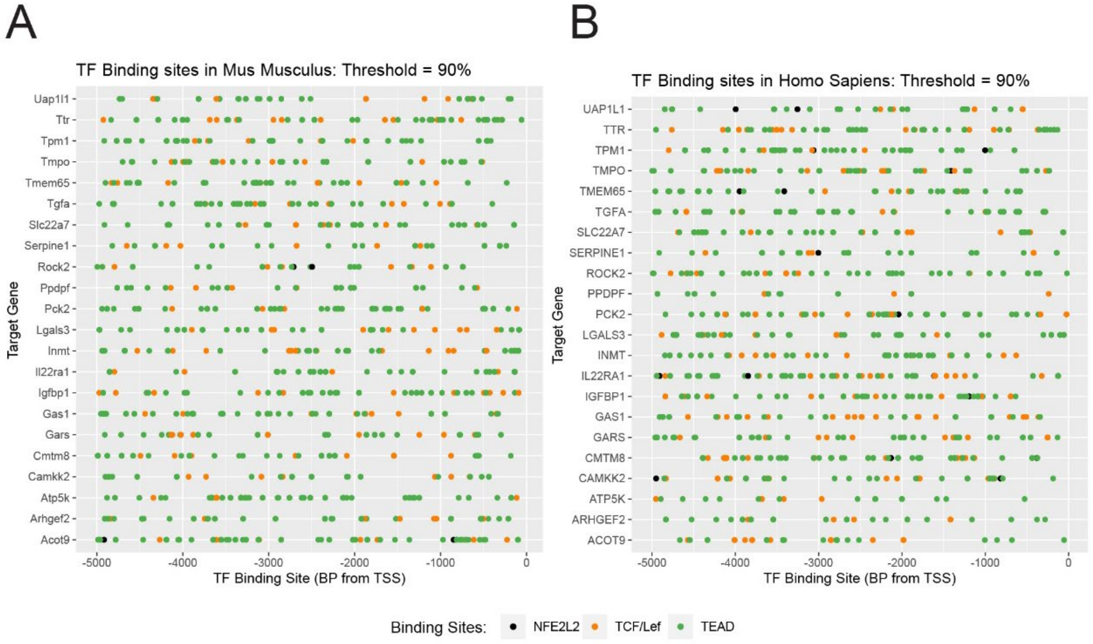


**Supplementary Figure S8.**
*In silico* promoter analysis. 5 kb of upstream promoter sequence for each of the genes listed in Table 1 were screened for the presence of consensus elements for Tcf/Lef, TEAD and ARE binding sites. **A,** murine genes. **B,** human genes.

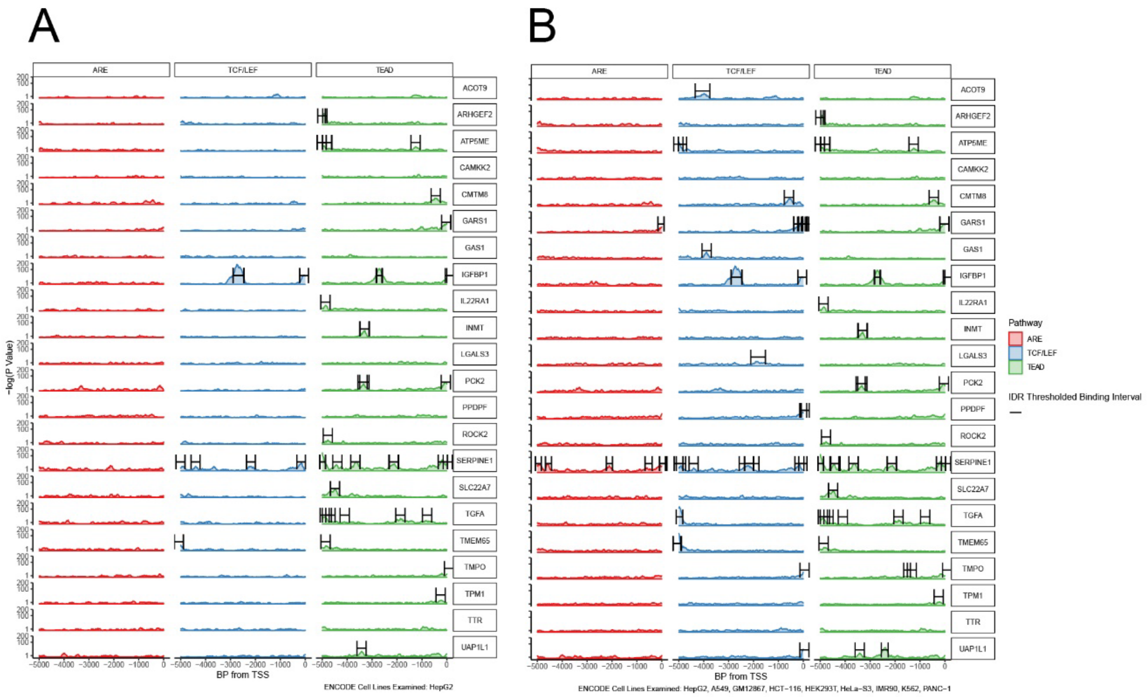

**Supplementary Figure S9.**
CHIP-seq results. **A,** True binding sites for Tcf/Lef, TEAD and ARE obtained from human HepG2 hepatoblastoma cells in the Encode v.5 data base (https://www.encodeproject.org/data-standards/chip-seq/) were mapped to the promoter sequences from Supplementary Fig. S8. **B,** Binding sites for Tcf/Lef, TEAD and ARE following the evaluation of Chip-Seq data from eight other human cell lines (see Materials and Methods).

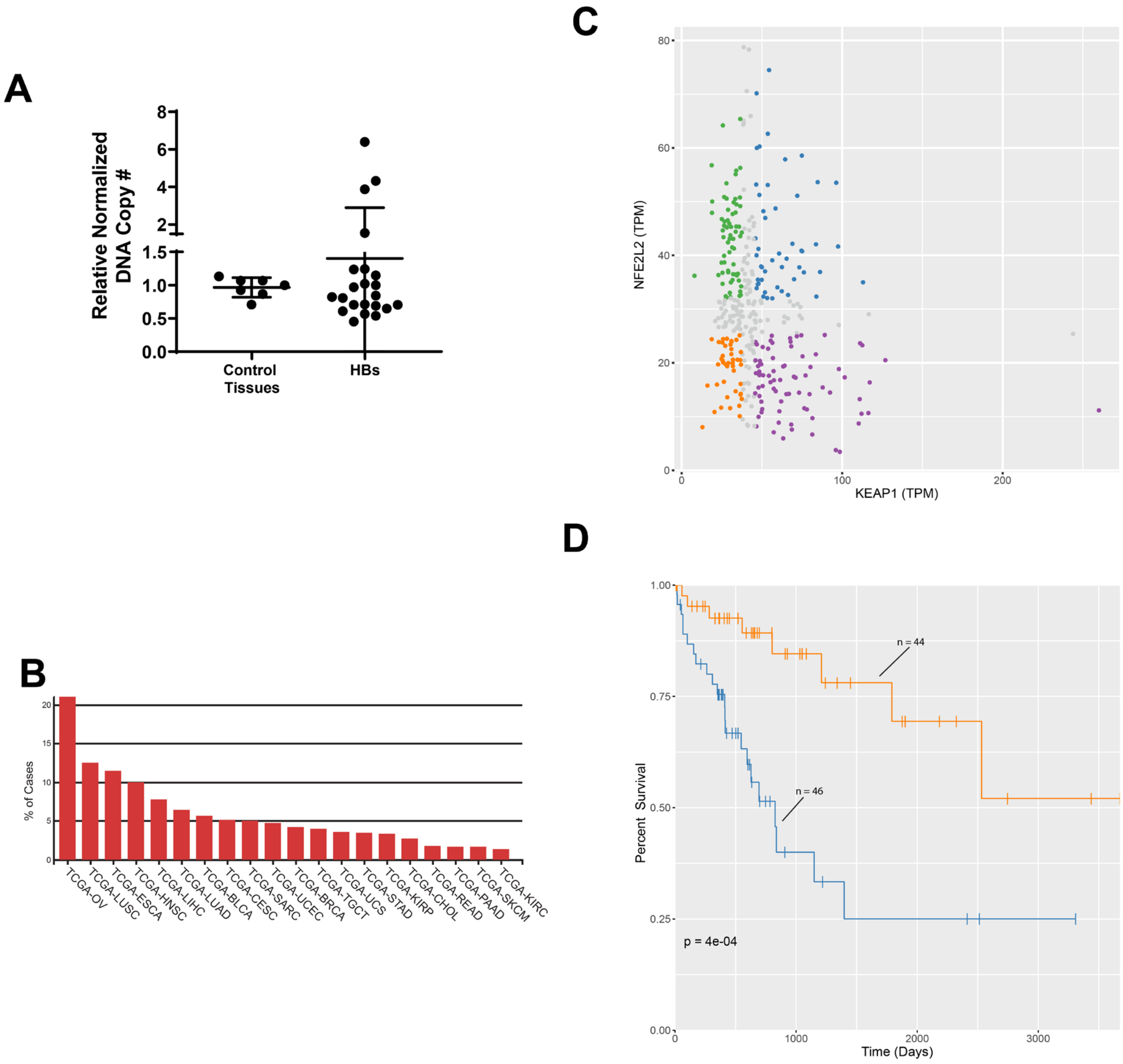


**Supplementary Figure S10.**

CNVs in HB and select human tumors. **A,** NFE2L2 CNVs determined from FFPE samples of seven normal tissues and 22 primary HBs. A TaqMan-based assay was used to quantify the copy number ratios of NFE2L2 versus two control genes (GAPDH and RPPH1). Each reaction was performed in triplicate and each point represents the calculated mean of the empirically-determined CNVs relative to those of the two control genes. **B,** Primary human tumors with the highest frequency of NFE2L2 gene amplification. Data were obtained from the Genomic Data Commons site in TCGA. **C,** Scatterplots of NFE2L2 and KEAP1 transcript expression in 371 HCC samples from the TCGA PANCAN data set. Sets are divided into quadrants that indicate those samples with the highest and lowest expression of NFE2L2 and KEAP1. **D,** Kaplan-Meier survival of individuals from **C** whose tumors contained the highest and lowest levels of NFE2L2 and KEAP1 expression.

### Supplementary Tables

#### Supplementary Table S1 : Top IPA Pathways Deregulated in HBs Expressing WT NFE2L2 vs. L30P/R34P

| Name of IPA Pathway | $\Delta$90+YAP$^{S127A}$ +L30P/R34P vs. $\Delta$90+YAP$^{S127A}$ Z-score | Name of IPA Pathway | $\Delta$90+YAP$^{S127A}$ +WT-NFE2L2 vs. $\Delta$90+YAP$^{S127A}$ Z-score | Name of IPA Pathway | $\Delta$90+YAP$^{S127A}$ +L30P/R34P vs. $\Delta$90+YAP$^{S127A}$ +WT-NFE2L2 Z-score |
|---|---|---|---|---|---|
| NRF2-mediated Oxidative Stress Response* | 3.888 | Oxidative Phosphorylation | -4.359 | Fcγ Receptor-mediated Phagocytosis in Macrophages and Monocytes | 3.71 |
| tRNA Charging | 3.771 | EIF2 Signaling | -3.128 | NF-κB Signaling | 3.266 |
| Leukocyte Extravasation Signaling | 3.43 | Sirtuin Signaling Pathway | 2.985 | Leukocyte Extravasation Signaling | 3.157 |
| Glutathione-mediated Detoxification* | 3.317 | NRF2-mediated Oxidative Stress Response* | 2.828 | IL-8 Signaling | 3 |
| Role of Pattern Recognition Receptors in Recognition of Bacteria and Viruses | 3 | | | Superpathway of Cholesterol Biosynthesis | -3 |
| Ephrin Receptor Signaling | 3 | | | Role of NFAT in Regulation of the Immune Response | 2.858 |
| RAN Signaling | 3 | | | Reelin Signaling in Neurons | 2.84 |
| Apelin Endothelial Signaling Pathway | 2.985 | | | LXR/RXR Activation | -2.828 |
| Fcγ Receptor-mediated Phagocytosis in Macrophages and Monocytes | 2.982 | | | fMLP Signaling in Neutrophils | 2.673 |
| Opioid Signaling Pathway | 2.92 | | | Fc Epsilon RI Signaling | 2.668 |
| fMLP Signaling in Neutrophils | 2.828 | | | NRF2-mediated Oxidative Stress Response* | 2.558 |
| Glutathione Redox Reactions I* | 2.828 | | | Apelin Endothelial Signaling Pathway | 2.53 |
| | | | | TREM1 Signaling | 2.53 |
| | | | | ERK5 Signaling | 2.53 |
| | | | | Role of Pattern Recognition Receptors in Recognition of Bacteria and Viruses | 2.496 |
| | | | | NER Pathway | 2.449 |
| | | | | Glutathione Redox Reactions I* | 2.449 |

* Red Indicates redox-regulated pathway

**Supplementary Table S2: The 821 gene expression differences shown in Fig. 4D and E Gene names and accession numbers are arranged in the order depicted in the Figure and as determined by hierarchical clustering. All values are expressed as CPM.**

| NAME | Δ90+YAP$^{S127A}$ | Δ90+YAP$^{S127A}$ +NFE2L2-WT | Δ90+YAP$^{S127A}$ +L30P/R34P |
|---|---|---|---|
| Lypla1 | 89.45 | 139.74 | 152.8 |
| Xkr9 | 4.33 | 11.73 | 18.12 |
| Smap1 | 36.67 | 50.95 | 59.67 |
| Tpp2 | 38.63 | 63.87 | 73.55 |
| Atic | 32.53 | 49.55 | 84.25 |
| Cab39 | 84.41 | 120.51 | 132.51 |
| Ugt1a6a | 62.01 | 117.42 | 211.19 |
| Ppip5k2 | 67.35 | 105.38 | 110.39 |
| Nr5a2 | 57.21 | 87.69 | 91.31 |
| BC003331 | 29.42 | 43.99 | 52.76 |
| Fmo4 | 7.12 | 15.2 | 15.71 |
| Degs1 | 89.35 | 132.95 | 193.21 |
| Lamc3 | 2.4 | 16.13 | 69.32 |
| Psmd5 | 31.64 | 45.49 | 66.86 |
| Stom | 36.59 | 79.26 | 100.1 |
| Ppp6c | 34.27 | 48.35 | 53.63 |
| Gtdc1 | 8.6 | 13.09 | 13.2 |
| Ifih1 | 24.47 | 35.97 | 39.96 |
| Qser1 | 28.19 | 46.93 | 51.06 |
| Cops2 | 47.59 | 68.7 | 79.07 |
| Srxn1 | 47.54 | 256.41 | 1463.35 |
| Tm9sf4 | 67.47 | 93.14 | 93.94 |
| Slc7a11 | 2.31 | 60.89 | 393.48 |
| Lxn | 1.66 | 8.89 | 13.43 |
| Arfip1 | 27.11 | 37.52 | 46.67 |
| Gba | 23.89 | 33.06 | 37.24 |
| S100a8 | 1.93 | 12.98 | 17.55 |
| S100a9 | 2.53 | 21.29 | 29.7 |
| Prune | 21.97 | 39.23 | 53.46 |
| Rnf115 | 26.29 | 36.9 | 50.3 |
| Trim33 | 25.17 | 36.15 | 36.96 |
| Chil3 | 0.47 | 5.59 | 7.49 |
| Cept1 | 36.43 | 57.59 | 71.67 |
| Gstm3 | 76.19 | 423.89 | 1508.79 |
| Gstm1 | 1335.19 | 3516.03 | 9635.56 |
| Gstm4 | 23.8 | 78.17 | 179.08 |
| Slc25a24 | 53.65 | 90.13 | 95.29 |
| Ctbs | 10.14 | 17.82 | 18.38 |
| Fpgt | 11.42 | 19.92 | 22.1 |
| Trp53inp1 | 74.92 | 116.68 | 155.1 |
| Cpne3 | 80.16 | 127.53 | 139.34 |
| Manea | 14.78 | 25.7 | 30.04 |
| Prkaa2 | 34.12 | 61.97 | 133.86 |
| Alpl | 14.4 | 30.7 | 35.79 |
| Cd36 | 67.48 | 177.93 | 237.38 |
| Bre | 23.82 | 37.24 | 40.3 |
| Cpeb2 | 22.91 | 36.71 | 39.19 |
| Pi4k2b | 28.74 | 40.69 | 42.04 |
| Ugdh | 113.66 | 275.17 | 842.9 |
| Dcun1d4 | 26.27 | 40.18 | 47.23 |
| Ugt2b34 | 340.34 | 703.37 | 1110.88 |
| Slc26a1 | 102.57 | 182.48 | 190.6 |
| Serpine1 | 11.66 | 31.64 | 67.88 |
| Met | 56.75 | 92.77 | 109.51 |
| Tmem209 | 19.11 | 28.18 | 36.89 |
| Akr1d1 | 105.66 | 207.32 | 212.13 |
| D630045J12Rik | 14.45 | 24.71 | 53.55 |
| Stambp | 12.3 | 19.38 | 21.48 |
| Eefsec | 23.19 | 34.05 | 38.43 |
| Styk1 | 0.2 | 5.47 | 19.35 |
| Cyp2a12 | 198.15 | 425.32 | 480.92 |
| Rhpn2 | 20.84 | 33.23 | 43.89 |
| AI987944 | 13.19 | 20.98 | 27.47 |
| C230091D08Rik | 7.19 | 12.45 | 13.5 |
| Mtmr10 | 32.26 | 45 | 46.26 |
| Picalm | 98.35 | 160.2 | 186.7 |
| Tsku | 51.69 | 132.28 | 315.24 |
| Tpp1 | 78.53 | 130.3 | 131.03 |
| Ppfibp2 | 14.53 | 24.14 | 55.59 |
| Acsm3 | 32.75 | 72.79 | 77.88 |
| Aldoa | 123.75 | 213.14 | 325.82 |
| Ifitm6 | 0.15 | 0.79 | 2.29 |
| Hpgd | 89.6 | 145.96 | 187.11 |
| Itfg1 | 49.12 | 71.11 | 90.99 |
| Ces2c | 2.65 | 13.95 | 15.51 |

#### Supplementary Table S2 (continued)

| NAME | Δ90+YAP$^{S127A}$ | Δ90+YAP$^{S127A}$ +NFE2L2-WT | Δ90+YAP$^{S127A}$ +L30P/R34P |
|---|---|---|---|
| Nfatc3 | 35.46 | 53.42 | 54.09 |
| Nqo1 | 6.7 | 37.31 | 222.5 |
| Bco1 | 13.87 | 26.88 | 43.97 |
| Mmp8 | 0.11 | 3.04 | 4.5 |
| Raver1 | 37.48 | 59.41 | 67.32 |
| Gclc | 169.37 | 522.46 | 2071.87 |
| Gsta1 | 0.86 | 72.14 | 1057.15 |
| Gm10639 | 0.08 | 6.46 | 136.35 |
| Gsta2 | 3.85 | 59.54 | 602.85 |
| Ngp | 1.35 | 16.06 | 21.3 |
| Ltf | 0.75 | 12.15 | 18.71 |
| Nt5dc1 | 9.17 | 14.56 | 18.9 |
| Reep3 | 75.77 | 112.21 | 166.78 |
| Ugp2 | 163.27 | 338.39 | 929.4 |
| Map2k4 | 31.41 | 51.24 | 51.38 |
| Srr | 31.71 | 54.73 | 60.77 |
| Taok1 | 58.63 | 87.55 | 87.83 |
| 5730455P16Rik | 13.75 | 19.72 | 21.55 |
| Usp32 | 35.65 | 53.59 | 55.03 |
| Mmd | 106.95 | 171.02 | 176.54 |
| Tom1l1 | 18.84 | 28.64 | 44.36 |
| Abcc3 | 94.89 | 194.96 | 341.12 |
| Pdk2 | 51.3 | 90.66 | 93.16 |
| Tmem106a | 13.81 | 22.34 | 25.14 |
| 1810010H24Rik | 8.47 | 15.5 | 24.47 |
| Stxbp6 | 24.32 | 34.92 | 36.3 |
| Egln3 | 18.42 | 32.52 | 41.74 |
| Cfl2 | 42.49 | 59.63 | 79.43 |
| Hif1a | 37.65 | 58.51 | 65.43 |
| Gpx2 | 0.72 | 7.79 | 60.66 |
| Susd6 | 58.33 | 85.04 | 91.57 |
| Gcnt2 | 19.6 | 30.89 | 34.47 |
| Ranbp9 | 28.6 | 39.73 | 41.18 |
| Ercc6l2 | 16.86 | 24.3 | 28.78 |
| Erap1 | 64.72 | 103.66 | 107.73 |
| Pxk | 28.69 | 40.06 | 47.74 |
| Vcl | 54.81 | 93.53 | 156.26 |
| Ap3m1 | 45.35 | 75.29 | 75.71 |
| Samd8 | 31.96 | 48.54 | 65.49 |
| Ppif | 24.28 | 36.59 | 37.13 |
| Ankrd28 | 36.35 | 56.63 | 61.64 |
| Ero1l | 26.48 | 43.63 | 61.13 |
| Abcc4 | 21.18 | 143.7 | 1011.15 |
| AW549877 | 29.84 | 46.89 | 47.71 |
| Golph3 | 64.21 | 98.32 | 103.56 |
| Laptm4b | 27.93 | 40.99 | 45.28 |
| Grina | 108.85 | 220.91 | 280.46 |
| Xpnpep3 | 21.22 | 30.07 | 31.2 |
| Slc38a2 | 117.97 | 180.55 | 211.31 |
| Dazap2 | 114.85 | 163.34 | 193.57 |
| Igfbp6 | 2.01 | 30.25 | 38.94 |
| Mapk1 | 84.35 | 121.93 | 145 |
| Cldn1 | 72.22 | 137.78 | 176.29 |
| Kpna1 | 34.79 | 53.14 | 64.03 |
| Pla1a | 22.73 | 43.61 | 63.2 |
| Retnlg | 0.2 | 2.87 | 3.76 |
| Tfg | 55.62 | 81.2 | 87.02 |
| Gbe1 | 51.72 | 114.83 | 266.29 |
| Mpc1 | 83.83 | 145.93 | 206.33 |
| BC004004 | 50.05 | 75.05 | 83.42 |
| Tbc1d5 | 21.81 | 32.19 | 37.54 |
| Sos1 | 26.21 | 37.38 | 38.3 |
| Lrpprc | 48.54 | 70.82 | 101.65 |
| Pcdhgc3 | 0.73 | 7.49 | 9.51 |
| Prr16 | 8.01 | 13.44 | 14.41 |
| Seh1l | 44.38 | 63.94 | 69.68 |
| Dym | 23.73 | 33.71 | 38.18 |
| Rtn3 | 111.43 | 165.12 | 213.03 |
| Asah2 | 18.33 | 28.73 | 31.76 |
| Ide | 76.06 | 123.18 | 152.95 |
| Gm10768 | 100.05 | 220.79 | 301.4 |
| Zdhhc6 | 44.81 | 63.13 | 65.56 |
| Maoa | 18.45 | 61.96 | 354.83 |
| Maob | 57.96 | 97.14 | 99.05 |
| Klhl13 | 20.78 | 41.31 | 87.91 |

#### Supplementary Table S2 (continued)

| NAME | Δ90+YAP$^{S127A}$ | Δ90+YAP$^{S127A}$ +NFE2L2-WT | Δ90+YAP$^{S127A}$ +L30P/R34P |
|---|---|---|---|
| Mtmr1 | 21.12 | 34.15 | 41.81 |
| Atp6ap1 | 71.45 | 99.48 | 102.03 |
| Ubqln2 | 28.95 | 46.68 | 58.58 |
| Rgl1 | 15.74 | 7.87 | 21.5 |
| Tor3a | 16.76 | 9 | 20.11 |
| Rcsd1 | 9.86 | 4.7 | 11.6 |
| Arhgap30 | 14.12 | 6.51 | 17.09 |
| Ifi203 | 7.17 | 3.55 | 9.19 |
| Tnfaip8l2 | 5.17 | 2.3 | 5.61 |
| Tinagl1 | 47.44 | 25.75 | 85.1 |
| Man1c1 | 10.19 | 4.53 | 10.76 |
| C1qc | 93.6 | 34 | 109.61 |
| C1qa | 126.92 | 51.33 | 146.21 |
| Anxa3 | 89.59 | 48.98 | 163.62 |
| Dok1 | 7.43 | 3.49 | 7.54 |
| Gmfg | 5.85 | 2.85 | 6.02 |
| Tmem86a | 14.88 | 7.31 | 17.3 |
| Lyve1 | 5.1 | 1.27 | 11.61 |
| Far1 | 12.87 | 4.44 | 13.58 |
| Pycard | 3.27 | 1.23 | 3.35 |
| Ifitm1 | 2.57 | 0.84 | 2.58 |
| Adgre5 | 35.42 | 19.07 | 49.96 |
| AB124611 | 4.58 | 1.85 | 8.1 |
| 2810417H13Rik | 13.86 | 6.3 | 14 |
| Cnn2 | 26.89 | 12.8 | 29.11 |
| Arhgap9 | 7.25 | 2.84 | 7.37 |
| Pmp22 | 17.51 | 4.3 | 23.63 |
| Slfn5 | 21.07 | 9.03 | 23 |
| Abi3 | 14.27 | 8.03 | 15.77 |
| Trib2 | 7.35 | 2.8 | 8.25 |
| Ly86 | 6.85 | 2.87 | 8.66 |
| Tgfbi | 50.18 | 24.18 | 52.2 |
| 4930486L24Rik | 4.61 | 0.85 | 8.44 |
| Ctla2b | 3.93 | 1.5 | 3.99 |
| Klhl6 | 5.7 | 2.52 | 6.27 |
| Hcls1 | 18.7 | 9.26 | 21.74 |
| Aif1 | 9.9 | 3.53 | 10.2 |
| Stard8 | 9.13 | 4.25 | 10.4 |
| Pcmtd1 | 54.33 | 84.1 | 82.95 |
| Tmem14a | 3.22 | 6.24 | 4.64 |
| Raph1 | 84.98 | 125.24 | 90.89 |
| Steap3 | 45.54 | 79.61 | 63.63 |
| Cdk18 | 30.87 | 48.61 | 41.59 |
| Cfhr3 | 2.49 | 6.29 | 4.39 |
| Gm4788 | 88.07 | 157.46 | 109.61 |
| Cfhr2 | 55.5 | 116.83 | 75.66 |
| Rabgap1l | 24.84 | 35.48 | 32.55 |
| Klhdc9 | 0.94 | 2.16 | 1.68 |
| Chml | 8.78 | 16 | 14.8 |
| Pter | 68.33 | 111.53 | 98.27 |
| Rsu1 | 30.76 | 45.51 | 38.79 |
| Stam | 19.08 | 28.52 | 26.5 |
| Mrps2 | 23.65 | 35.93 | 34.65 |
| 1110008P14Rik | 12.85 | 19.73 | 14.42 |
| Nr6a1 | 5.94 | 10.38 | 6.79 |
| Acvr2a | 25.66 | 38.05 | 33.01 |
| Nfe2l2 | 140.83 | 391.99 | 258.8 |
| Zfp385b | 15.93 | 26.34 | 22.23 |
| Tcp11l1 | 13.76 | 23.85 | 19.35 |
| BC052040 | 8.67 | 13.65 | 13.56 |
| Adal | 12.04 | 17.5 | 17 |
| Psmf1 | 27.15 | 42.98 | 41.44 |
| Ncoa6 | 34.45 | 54.35 | 47.24 |
| Rbm38 | 5.08 | 9.74 | 8.59 |
| Bche | 67.8 | 139.57 | 118.33 |
| Sf3b4 | 28.73 | 39.36 | 37.84 |
| Hfe2 | 27.45 | 70.4 | 52.96 |
| Gdap2 | 32.81 | 51.17 | 47.04 |
| Alg14 | 10.85 | 18.5 | 15.22 |
| Sec24b | 40 | 55.41 | 55.08 |
| Ppa2 | 26.43 | 37.65 | 30.54 |
| Mob3b | 31.1 | 56.08 | 52.48 |
| Angptl3 | 199.22 | 420.99 | 396.87 |
| Tmem69 | 10.57 | 16.15 | 13.97 |
| Elovl1 | 53.65 | 87.06 | 80.61 |

#### Supplementary Table S2 (continued)

| NAME | Δ90+YAP$^{S127A}$ | Δ90+YAP$^{S127A}$ +NFE2L2-WT | Δ90+YAP$^{S127A}$ +L30P/R34P |
|---|---|---|---|
| Mapre3 | 20.49 | 38.24 | 36.57 |
| Atxn2 | 50.53 | 72.82 | 66.9 |
| Orai1 | 17.32 | 25.14 | 24.63 |
| Cyp3a13 | 27.17 | 52.43 | 35.02 |
| Bri3 | 35.06 | 58.18 | 51.21 |
| Baiap2l1 | 37.99 | 60.93 | 55.46 |
| Cyp3a59 | 9.64 | 25.11 | 22.22 |
| Peg10 | 3.72 | 10.97 | 4.56 |
| Wasl | 74.9 | 115.12 | 114.47 |
| Zfp800 | 11.76 | 17.99 | 17.31 |
| Mkln1 | 45.36 | 63.32 | 60.26 |
| Zyx | 49.31 | 79.37 | 63.65 |
| Wipf3 | 26.75 | 47.06 | 37.94 |
| Kdm3a | 19.14 | 29.59 | 25.73 |
| Tmsb10 | 44.91 | 89.91 | 83.42 |
| Frmd4b | 51.64 | 79.66 | 67.8 |
| Bhlhe40 | 81.57 | 149.58 | 116.64 |
| Setd5 | 61.83 | 87.7 | 78.67 |
| Tmcc1 | 20.31 | 31.2 | 27.61 |
| Slc6a12 | 31.37 | 57.03 | 32.13 |
| Atn1 | 54.43 | 81.56 | 67.41 |
| Ceacam1 | 60.26 | 119.53 | 71.5 |
| Zfp36 | 39.13 | 63.74 | 61.97 |
| Ankrd27 | 69.16 | 103.14 | 93.37 |
| Gm5595 | 2.34 | 4.38 | 3.37 |
| Kdelr1 | 83.51 | 124.55 | 113.25 |
| Cpeb1 | 17.31 | 29.78 | 23.28 |
| Tmem135 | 53.05 | 90.49 | 89.33 |
| Ints4 | 27.13 | 39 | 35.78 |
| Ppme1 | 29.34 | 44.16 | 42.96 |
| Gm5601 | 1.15 | 7.73 | 6.3 |
| Sult1a1 | 490.91 | 1021.73 | 852.73 |
| Cdhr5 | 80.31 | 126.58 | 80.57 |
| Slc22a18 | 53.09 | 109.69 | 103.71 |
| Rbpms | 67.37 | 108.18 | 98.15 |
| Mtus1 | 167.03 | 239.33 | 196.71 |
| Aadat | 19.52 | 37.21 | 34.78 |
| Tm6sf2 | 18.08 | 39.26 | 29.34 |
| Tom1 | 31.07 | 49.69 | 41.18 |
| Slc38a7 | 27.41 | 36.63 | 34.82 |
| D230025D16Rik | 37.03 | 57.92 | 53.25 |
| Acp5 | 59.61 | 117.36 | 114.73 |
| Vwa5a | 57.83 | 98.85 | 94.02 |
| Ubl7 | 22.45 | 36.93 | 34.54 |
| Dennd4a | 30.9 | 51.6 | 50.15 |
| Rbpms2 | 17.96 | 30.87 | 28.11 |
| Ick | 25.98 | 39.71 | 34.84 |
| Rn7sk | 2.29 | 8.44 | 5.72 |
| Abhd14b | 96.33 | 158.2 | 137.95 |
| Camp | 0.75 | 4.7 | 4.33 |
| Rtp3 | 30.82 | 47.08 | 30.97 |
| Cyp8b1 | 58.36 | 168.07 | 155.8 |
| Tcaim | 9.2 | 15.9 | 14.23 |
| Lars2 | 72.21 | 272.09 | 234.47 |
| Tab2 | 78.38 | 113.66 | 108.98 |
| Sgk1 | 17.78 | 37.97 | 34.7 |
| Ppa1 | 80.07 | 128.01 | 125.47 |
| Scyl2 | 39.22 | 60.79 | 59.7 |
| Mettl7b | 225.22 | 459.77 | 318.37 |
| Tug1 | 52.38 | 77.79 | 62.69 |
| Ube2d-ps | 4.66 | 8.08 | 6.14 |
| 0610010F05Rik | 16.88 | 24.42 | 24.12 |
| Timd2 | 41.81 | 80.15 | 76.21 |
| Sec24a | 65.07 | 103.41 | 96.32 |
| Drg2 | 28.75 | 40.64 | 39.45 |
| Asgr2 | 177.41 | 329.02 | 297.99 |
| Ggt6 | 13.19 | 26.11 | 15.77 |
| Ctns | 17.26 | 26.63 | 21.07 |
| Serpinf1 | 205.61 | 483.72 | 448.1 |
| Ccl9 | 25.18 | 52.84 | 33.98 |
| Dcakd | 37.13 | 51.4 | 41.7 |
| Prkca | 13 | 20.87 | 18.17 |
| Fam20a | 26.19 | 45.73 | 29.56 |
| Afmid | 61.55 | 112.06 | 96.22 |
| Gphn | 33.87 | 52.54 | 50.62 |

#### Supplementary Table S2 (continued)

| NAME | Δ90+YAP$^{S127A}$ | Δ90+YAP$^{S127A}$ +NFE2L2-WT | Δ90+YAP$^{S127A}$ +L30P/R34P |
|---|---|---|---|
| Numb | 31.03 | 43.67 | 36.51 |
| Serpina3n | 158.42 | 380.34 | 238.4 |
| Akr1e1 | 20.54 | 33.3 | 27.5 |
| Serpinb9 | 12.42 | 20.88 | 15.1 |
| Zfp935 | 3.5 | 6.21 | 5.57 |
| Zfp729b | 6.04 | 10.1 | 9.31 |
| Thrb | 23.55 | 35.75 | 31.54 |
| Parp4 | 28.74 | 46.73 | 44.43 |
| Gm17066 | 13.04 | 20.74 | 15.74 |
| Cpq | 59.01 | 84.48 | 68.31 |
| Rnf139 | 32.66 | 48.68 | 44.86 |
| St3gal1 | 83.01 | 146.19 | 119.02 |
| Pkp2 | 70.82 | 103.65 | 85.89 |
| Hrg | 204.21 | 569.6 | 423.3 |
| Kng1 | 1442.91 | 2828.61 | 1958.11 |
| Pcyt1a | 77.65 | 110.79 | 110.34 |
| Gm38396 | 12.19 | 19.71 | 17.02 |
| Zfp820 | 2.88 | 5.46 | 4.53 |
| Zfp942 | 7.91 | 13.58 | 12.12 |
| Zfp946 | 9.43 | 15.1 | 13.57 |
| Tbc1d24 | 25.91 | 43.77 | 35.17 |
| Ppt2 | 23.38 | 32.77 | 31.23 |
| Vmac | 15.66 | 22.64 | 16.81 |
| Slc25a23 | 179.08 | 307.72 | 230.01 |
| Galm | 46.62 | 73.36 | 69.25 |
| Fbxo11 | 29.4 | 41.07 | 40.87 |
| Wac | 55.63 | 81.53 | 77.35 |
| Dsc2 | 112.75 | 192.29 | 122.18 |
| Patl1 | 25.02 | 39.91 | 39.62 |
| Zdhhc9 | 78.54 | 141.52 | 131.2 |
| Mospd2 | 31.8 | 47.32 | 47.02 |
| Cox5b | 88.09 | 55.33 | 54.83 |
| Dtymk | 37.66 | 22.58 | 22.17 |
| Prrx1 | 11.68 | 3.42 | 1.92 |
| Mrpl41 | 26.04 | 14.14 | 13.63 |
| Tmem203 | 14.14 | 8.83 | 7.13 |
| Phpt1 | 37.18 | 14.95 | 13.4 |
| Adamtsl2 | 13.34 | 4.32 | 3.45 |
| Ak1 | 3.77 | 1.4 | 1.35 |
| Ndufa8 | 100.36 | 56.25 | 49.14 |
| Olfml2a | 3.33 | 0.96 | 0.6 |
| Ppig | 78.61 | 53.94 | 48.83 |
| Nop10 | 31.33 | 19.17 | 19.16 |
| Rtf1 | 53.42 | 38.64 | 37.61 |
| Serf2 | 307.71 | 188.33 | 150.22 |
| Hypk | 90.71 | 42.61 | 41.61 |
| Romo1 | 45.41 | 15.1 | 12.9 |
| D630003M21Rik | 22.93 | 7.39 | 6.64 |
| Pkig | 24.82 | 16.52 | 14.53 |
| Dnttip1 | 34.82 | 22.7 | 20.86 |
| Atp5e | 137.66 | 63.21 | 55.58 |
| Ndufc1 | 29.16 | 16.46 | 14.33 |
| Apoa1bp | 51.85 | 32.45 | 31.7 |
| Dpm3 | 23.38 | 9.25 | 8.27 |
| Mrps21 | 39.44 | 22.3 | 22.09 |
| Gm20752 | 9.58 | 2.13 | 0.96 |
| Chchd7 | 11.39 | 6.53 | 6.49 |
| Svbp | 14.56 | 7.28 | 7.13 |
| Ndufs5 | 11.31 | 4.34 | 3.95 |
| Smim12 | 23.12 | 15.28 | 14.57 |
| Rnf19b | 53.5 | 37.37 | 34.67 |
| Tmem234 | 78.74 | 55.77 | 54.8 |
| Minos1 | 65.6 | 37.63 | 34.75 |
| Pdpn | 4.65 | 0.63 | 0.33 |
| Gpr153 | 5.22 | 1.55 | 1.49 |
| Fam133b | 18.78 | 10.63 | 10.21 |
| Fzd1 | 19.59 | 10.04 | 7.6 |
| Reln | 42.8 | 15.39 | 10.6 |
| Mxd4 | 44.98 | 30 | 29.9 |
| Bloc1s4 | 17.43 | 10.72 | 9.98 |
| Igfbp7 | 357.64 | 174.93 | 142.72 |
| Mrps18c | 22.97 | 14.89 | 14.82 |
| BC005561 | 12.22 | 7.19 | 5.59 |
| Atp5k | 43.53 | 9.1 | 8.67 |
| Srsf9 | 48.61 | 31.14 | 31.01 |

#### Supplementary Table S2 (continued)

| NAME | Δ90+YAP$^{S127A}$ | Δ90+YAP$^{S127A}$ +NFE2L2-WT | Δ90+YAP$^{S127A}$ +L30P/R34P |
|---|---|---|---|
| Shfm1 | 59.7 | 32.95 | 29.99 |
| Ndufa5 | 44 | 22.83 | 22.73 |
| 1110001J03Rik | 17.26 | 6.01 | 4.24 |
| Ndufb2 | 42.87 | 18.78 | 18.6 |
| Tacstd2 | 5.93 | 0.45 | 0.16 |
| Vamp5 | 38.33 | 17.02 | 12.8 |
| Bola3 | 27.65 | 15.16 | 15.02 |
| Snrpg | 39.71 | 15.42 | 13.65 |
| Brk1 | 86.53 | 60.27 | 53.59 |
| Grcc10 | 84.33 | 41.3 | 36.24 |
| Lockd | 4.3 | 1.27 | 1.1 |
| H2afj | 79.1 | 38.24 | 26.52 |
| Mgp | 34.09 | 9.23 | 5.29 |
| Ndufa3 | 49.07 | 17.08 | 15.61 |
| Tfpt | 13.97 | 8.29 | 7.34 |
| Inafm1 | 14.77 | 7.87 | 6.77 |
| Snrpd2 | 61.37 | 31.35 | 28.73 |
| Rabac1 | 134.59 | 87.97 | 82.21 |
| Megf8 | 44.47 | 30.63 | 28.7 |
| Eid2 | 6.45 | 2.17 | 1.66 |
| Cox7a1 | 1.59 | 0.33 | 0.32 |
| Pdcd5 | 37.26 | 24.47 | 20.22 |
| Saa2 | 88.94 | 8.8 | 7.1 |
| Dkk3 | 9.33 | 2.88 | 2.53 |
| Hs3st2 | 16.95 | 2.11 | 1.69 |
| Bola2 | 26.13 | 11.95 | 10.29 |
| Gm4532 | 1.31 | 0.33 | 0.13 |
| Bcl7c | 42.93 | 26.35 | 19.93 |
| Bet1l | 25.73 | 16.9 | 15.55 |
| Pet100 | 11.56 | 3.93 | 3.47 |
| Arglu1 | 56.6 | 33.91 | 24.37 |
| Ing1 | 36.66 | 25.93 | 23.37 |
| Ndufa13 | 83.63 | 52.1 | 44.24 |
| Ccdc124 | 46.46 | 26.16 | 20.81 |
| Use1 | 64.31 | 41.96 | 37.98 |
| D8Ertd738e | 79.73 | 55.29 | 50.42 |
| 2310036O22Rik | 99.83 | 66.33 | 54.98 |
| Ccdc102a | 5.92 | 1.94 | 1.81 |
| Fam96b | 27.1 | 12.32 | 12.01 |
| Zfhx3 | 69.76 | 41.45 | 36.16 |
| Ubl5 | 61.43 | 39.29 | 34.42 |
| Tmed1 | 29.15 | 19.1 | 15.25 |
| Cib2 | 16.92 | 8.5 | 4.16 |
| Snapc5 | 24.35 | 14.03 | 11.99 |
| Anapc13 | 43.95 | 17.88 | 17.69 |
| Ccdc12 | 26.09 | 16.54 | 14.97 |
| Higd1a | 34.72 | 20.18 | 18.94 |
| A330049N07Rik | 11.68 | 2.51 | 1.38 |
| 2310011J03Rik | 35.93 | 23.27 | 21.74 |
| Lsm7 | 33.21 | 16.19 | 13.66 |
| Timm13 | 86.81 | 42.06 | 40.36 |
| Csrp2 | 115.33 | 76.28 | 65.79 |
| Arhgef25 | 4.71 | 2.11 | 1.89 |
| Cnpy2 | 91.25 | 56.58 | 46.61 |
| Cd63 | 167.02 | 99.57 | 84.42 |
| Selm | 5.3 | 2.22 | 1.66 |
| Mrps24 | 39.66 | 26.42 | 24.06 |
| H2afv | 73 | 46.85 | 38.81 |
| Hba-a2 | 11.52 | 0.95 | 0.31 |
| Leap2 | 12.08 | 4.06 | 3.29 |
| Atox1 | 55.27 | 31.38 | 30.41 |
| Dhrs7b | 28.81 | 18.76 | 18.18 |
| Chd3 | 83.13 | 54.45 | 47.21 |
| Tmem256 | 41.08 | 15.23 | 11.22 |
| Sdf2 | 49.08 | 34.19 | 31.53 |
| Supt4a | 45.51 | 27.06 | 23.87 |
| Cuedc1 | 21.71 | 10.4 | 8.25 |
| Cbx1 | 50.23 | 30.81 | 26.48 |
| P3h4 | 2.67 | 0.81 | 0.73 |
| Mrc2 | 13.56 | 3.66 | 3.4 |
| Ict1 | 43.68 | 28.45 | 25.86 |
| Foxj1 | 4.98 | 1.59 | 0.57 |
| Degs2 | 6.24 | 2.04 | 0.89 |
| Inhba | 37.3 | 16.11 | 8.02 |
| Sox4 | 118.36 | 43.03 | 21.15 |

#### Supplementary Table S2 (continued)

| NAME | Δ90+YAP$^{S127A}$ | Δ90+YAP$^{S127A}$ +NFE2L2-WT | Δ90+YAP$^{S127A}$ +L30P/R34P |
|---|---|---|---|
| Idnk | 22.87 | 15.34 | 11.45 |
| Med10 | 22.85 | 15.81 | 14.86 |
| Ndufs6 | 84.65 | 30.36 | 28.51 |
| Cetn3 | 46.2 | 31.81 | 31.63 |
| Smim4 | 7.21 | 2.86 | 2.44 |
| Gdf10 | 13.81 | 3.02 | 2.83 |
| Mmrn2 | 43.63 | 10.95 | 9.6 |
| Dpysl2 | 11.07 | 5.66 | 4.84 |
| Polr2k | 16.87 | 8.16 | 6.77 |
| Polr2f | 37.08 | 19.83 | 18.18 |
| Cbx6 | 66.82 | 39.18 | 24.35 |
| Rps19bp1 | 27.7 | 16.83 | 14.56 |
| Smdt1 | 65.94 | 44.12 | 36.07 |
| Creld2 | 130.09 | 46.17 | 43.86 |
| Cox14 | 43.62 | 22.36 | 21.61 |
| Smim22 | 2.67 | 0.9 | 0.64 |
| Scarf2 | 10.52 | 2.93 | 1.93 |
| Ndufb4 | 30.8 | 17.58 | 16.17 |
| Atp5j | 135.76 | 88.12 | 85.18 |
| Smim11 | 19.48 | 11.8 | 8.24 |
| Tmem242 | 52.22 | 36.06 | 33.14 |
| Gtf2h5 | 43.61 | 30.5 | 29.85 |
| Nme3 | 28.81 | 15.28 | 12.82 |
| Fam173a | 48.82 | 27.67 | 25.5 |
| Adamts10 | 18.49 | 9.82 | 4.99 |
| Rps18 | 21.23 | 11.81 | 10.01 |
| 1110038B12Rik | 25.33 | 14.1 | 13.53 |
| Gnl1 | 40.31 | 29.68 | 27.71 |
| Polr1c | 40.75 | 26.94 | 26.84 |
| Ptprs | 84.85 | 54.65 | 33.85 |
| Pura | 49.55 | 35.97 | 32.58 |
| Tmem134 | 58.06 | 39.35 | 35.99 |
| Ccs | 138.03 | 85.71 | 73.39 |
| Ccdc85b | 61.97 | 35.94 | 23.1 |
| Cdc42bpg | 78.91 | 48.71 | 46.07 |
| Trmt112 | 58.9 | 41.06 | 40.5 |
| Esrra | 35.96 | 22.5 | 21.97 |
| Bad | 28.17 | 16.61 | 14.42 |
| Ppp1r14b | 68.89 | 33.68 | 31.06 |
| Fkbp2 | 80.32 | 49.57 | 42.18 |
| 2700081O15Rik | 23.13 | 12.92 | 7.88 |
| Uqcc3 | 29.93 | 19.28 | 15.59 |
| Npm3 | 29.93 | 17.54 | 15.54 |
| Usmg5 | 68.19 | 31.02 | 28.77 |
| Pin4 | 17.98 | 9.76 | 9.59 |
| Rpl34 | 7.99 | 2.87 | 2.87 |
| Rab26os | 1.04 | 0.13 | 0.13 |
| Ptpn18 | 4.32 | 1.43 | 3.13 |
| Sema4c | 10.09 | 5.45 | 6.17 |
| Rpl31 | 377.92 | 190.37 | 202.11 |
| Fhl2 | 3.84 | 1.19 | 1.62 |
| Col3a1 | 159.2 | 52.26 | 107.44 |
| Hspe1 | 168.18 | 79.72 | 85.2 |
| Ndufb3 | 30.73 | 18.18 | 21.69 |
| Rpl37a | 330.55 | 144.64 | 158.05 |
| Myeov2 | 42.28 | 14.58 | 15.73 |
| Fam174a | 15.22 | 9.67 | 11.78 |
| Snrpe | 31.42 | 20.26 | 22.69 |
| Ppfia4 | 9.09 | 2.75 | 6.58 |
| Tmem9 | 35.9 | 24.07 | 25.44 |
| Rnasel | 10.24 | 3.54 | 7.26 |
| Pfdn2 | 35.06 | 18.3 | 19.84 |
| Ackr1 | 1.16 | 0.31 | 0.77 |
| Itpkb | 15.46 | 6.4 | 11.26 |
| Hlx | 6.88 | 2.34 | 6.84 |
| Batf3 | 1.7 | 0.47 | 1 |
| Fam171a1 | 5.11 | 2.28 | 3.24 |
| Celf2 | 23.14 | 12.67 | 18.18 |
| Itga8 | 7.81 | 2.64 | 4.77 |
| Zmynd19 | 21.18 | 13.92 | 16.14 |
| Egfl7 | 22.7 | 11.29 | 13.83 |
| Eng | 51.74 | 26.89 | 45.09 |
| Rpl35 | 304.07 | 167.77 | 202.38 |
| Klhl23 | 5.78 | 2.03 | 2.17 |
| 2700094K13Rik | 19.42 | 11.56 | 15.57 |

#### Supplementary Table S2 (continued)

| NAME | Δ90+YAP$^{S127A}$ | Δ90+YAP$^{S127A}$ +NFE2L2-WT | Δ90+YAP$^{S127A}$ +L30P/R34P |
|---|---|---|---|
| Gm13889 | 4.73 | 1.39 | 2.07 |
| Ccdc34 | 23.59 | 12.21 | 13.93 |
| Fbn1 | 18.9 | 6.02 | 9.79 |
| 1500011K16Rik | 18.11 | 8.01 | 8.66 |
| Mrps26 | 35.04 | 22.33 | 24.57 |
| Prnd | 4.74 | 0.76 | 1.43 |
| Gm561 | 7.3 | 3.44 | 4.23 |
| Top1 | 85.15 | 58.31 | 59.99 |
| Mybl2 | 6.07 | 2.51 | 3.83 |
| Zfas1 | 31.29 | 8.04 | 11.57 |
| Tshz2 | 29.38 | 9.74 | 10.04 |
| Rps21 | 264.86 | 83.27 | 99.66 |
| Rpl22l1 | 83.55 | 32.28 | 42.78 |
| Lhfp | 17.92 | 6.51 | 7.17 |
| Smc4 | 78.88 | 46.44 | 70.11 |
| Gucy1a3 | 9.28 | 3.79 | 5.07 |
| Nes | 13.98 | 5.09 | 5.98 |
| Rps27 | 288.21 | 125.96 | 141.01 |
| Mrpl9 | 44.34 | 30.17 | 33.37 |
| Ctsk | 3.06 | 0.74 | 0.79 |
| Olfml3 | 9.59 | 3.47 | 4.33 |
| Lamtor5 | 31.67 | 22.46 | 23.97 |
| Snhg8 | 8.68 | 3.5 | 3.53 |
| Gar1 | 21.13 | 11.22 | 13.12 |
| H2afz | 116.01 | 59.48 | 87.7 |
| Clca3a1 | 16.55 | 7.31 | 9.22 |
| Ndufb6 | 49.86 | 23.35 | 30.32 |
| Tomm5 | 24.13 | 14.1 | 18.98 |
| Rgs3 | 15.01 | 6.24 | 8.26 |
| Tm2d1 | 16.34 | 10.96 | 13.13 |
| Tie1 | 18.19 | 7.08 | 11.52 |
| Heyl | 6.49 | 1.82 | 2.53 |
| 1110065P20Rik | 24.6 | 13.44 | 13.76 |
| Eva1b | 9.67 | 3.55 | 4.28 |
| Trappc3 | 41.52 | 26.67 | 27.75 |
| Gja4 | 5.81 | 1.3 | 2.41 |
| Hpca | 1.9 | 0.12 | 0.14 |
| Hdac1 | 30.59 | 21.46 | 22.9 |
| Fam167b | 6.81 | 1.68 | 3.13 |
| Med18 | 5.15 | 2.36 | 3.42 |
| Hmgn2 | 67.67 | 27.93 | 28.93 |
| Stmn1 | 84.74 | 42.75 | 56.18 |
| Rpl11 | 435.33 | 242.69 | 289.01 |
| Gm13056 | 0.96 | 0.13 | 0.37 |
| Rpl22 | 193.59 | 117.63 | 146.4 |
| Tomm7 | 31.27 | 11.7 | 12.3 |
| Nos3 | 4.76 | 2.34 | 3.58 |
| Cenpa | 18.64 | 9.77 | 13.91 |
| Ost4 | 69.53 | 45.57 | 45.81 |
| Emilin1 | 55.28 | 14.15 | 23.45 |
| Sorcs2 | 8.26 | 0.33 | 0.47 |
| Med28 | 29.91 | 21.48 | 21.74 |
| Rpl9 | 413.81 | 232.36 | 296.61 |
| Plac8 | 11.12 | 3.84 | 10.24 |
| 1500011B03Rik | 4.61 | 2.02 | 3.05 |
| Pebp1 | 268.37 | 169.31 | 180.42 |
| Snrnp35 | 7.72 | 4.03 | 4.39 |
| Eln | 19.24 | 5.22 | 5.72 |
| Pdap1 | 91.05 | 60.23 | 66.68 |
| Rpa3 | 11.28 | 6.64 | 6.72 |
| Lsm8 | 12.11 | 7.32 | 8.56 |
| Gimap6 | 9.08 | 3.44 | 6.55 |
| Aqp1 | 53.56 | 20.64 | 46.62 |
| Lsm5 | 8.8 | 4.77 | 5 |
| Dysf | 13.19 | 4.53 | 12.32 |
| Fgd5 | 7.15 | 3.28 | 5.24 |
| Rpl32 | 400.38 | 215.5 | 270.63 |
| Plxnd1 | 60.17 | 28.66 | 48.25 |
| Gm8203 | 61.79 | 29.32 | 52.98 |
| P3h3 | 7.31 | 2.61 | 3.11 |
| Ccnd2 | 24.47 | 9.78 | 12.75 |
| Rerg | 2.99 | 1.01 | 1.47 |
| U2af2 | 120.04 | 85.45 | 89.95 |
| Tmem160 | 36.31 | 21.27 | 22.09 |
| Exoc3l2 | 7.96 | 2.31 | 3.98 |

**Supplementary Table S2 (continued)**

| NAME | Δ90+YAP$^{S127A}$ | Δ90+YAP$^{S127A}$ +NFE2L2-WT | Δ90+YAP$^{S127A}$ +L30P/R34P |
|---|---|---|---|
| Rps19 | 372.35 | 184.52 | 215.17 |
| Ltbp4 | 35.35 | 13.66 | 16.95 |
| Rps16 | 505.76 | 235.39 | 295.09 |
| Polr2i | 15.12 | 9.53 | 11.56 |
| Tyrobp | 27.79 | 11.38 | 25.73 |
| Clec11a | 2.54 | 0.94 | 1.04 |
| Rcn3 | 9.4 | 2.38 | 4.75 |
| Rps11 | 425.19 | 231 | 270.31 |
| Rpl13a | 1089.24 | 532.05 | 576.18 |
| Rpl18 | 473.29 | 251.45 | 304.27 |
| Chsy1 | 8.41 | 4.07 | 5.99 |
| Igf1r | 30.47 | 6.07 | 10.6 |
| Rps17 | 334.86 | 178.01 | 214.05 |
| Ndufc2 | 47.01 | 32.14 | 32.37 |
| Serpinh1 | 48.5 | 17.63 | 20.7 |
| Rps3 | 602.17 | 330.12 | 415.74 |
| Tmem159 | 8.8 | 4.37 | 4.99 |
| 4930413G21Rik | 5.52 | 2.47 | 2.61 |
| Hirip3 | 13.03 | 7.15 | 10.73 |
| Ctbp2 | 8.47 | 2.97 | 4.62 |
| Rplp2 | 273.98 | 165.52 | 189.24 |
| Tssc4 | 28.95 | 16.58 | 19.36 |
| Osbpl5 | 4.65 | 2.31 | 3.24 |
| Gins4 | 23.07 | 16.55 | 18.16 |
| Adgra2 | 11.21 | 3.44 | 6.73 |
| Hand2 | 3.71 | 0.97 | 2.07 |
| Hmgb2 | 55.2 | 25.67 | 41.42 |
| Nr2c2ap | 23.28 | 15.57 | 17.68 |
| Lsm4 | 38.06 | 19.61 | 22.38 |
| Mpv17l2 | 22.06 | 14.78 | 14.91 |
| Ushbp1 | 10.72 | 4.98 | 5.23 |
| Ankle1 | 3.79 | 1.38 | 2.49 |
| Plvap | 51.69 | 21.56 | 42.93 |
| Cklf | 2.57 | 1.12 | 1.14 |
| Exoc3l | 3.89 | 1.54 | 2.68 |
| Glg1 | 76.72 | 48.08 | 59.13 |
| Maf | 32.39 | 10.66 | 30.09 |
| Hsbp1 | 101.57 | 67.64 | 70.02 |
| Gins2 | 10.96 | 4.72 | 10.6 |
| Irf8 | 24.14 | 10.06 | 20.69 |
| Cyba | 23.48 | 9.73 | 20.5 |
| Amotl1 | 30.37 | 6.6 | 8.49 |
| Pin1 | 37.53 | 21.23 | 23.21 |
| Cdkn2d | 9.33 | 5.36 | 5.49 |
| Spc24 | 15.43 | 7 | 10.21 |
| Gm10698 | 41.24 | 13.39 | 21.2 |
| Esam | 21.02 | 9.57 | 12.75 |
| Il10ra | 12.28 | 6.07 | 12.27 |
| Tagln | 11.61 | 3.82 | 6.46 |
| Rps27l | 155.95 | 96.54 | 101.68 |
| Chst2 | 11.57 | 2.71 | 4.2 |
| Itga9 | 34.84 | 17.8 | 31.52 |
| Vipr1 | 10.94 | 3.12 | 6.31 |
| Fbxo5 | 5.84 | 2.52 | 4.3 |
| Tcf21 | 2.21 | 0.57 | 1.05 |
| Ctgf | 78.78 | 39.39 | 41.1 |
| Rspo3 | 2.58 | 0.6 | 1.18 |
| Cenpw | 3.61 | 1.14 | 1.68 |
| Hint3 | 18.58 | 11.61 | 12.39 |
| Fyn | 9.83 | 4.63 | 8.24 |
| Snrpd3 | 52.32 | 28.76 | 35.36 |
| Derl3 | 22.35 | 3.98 | 6.94 |
| Oaz1 | 492.97 | 249.73 | 322.34 |
| Arl1 | 66.35 | 44.04 | 48.73 |
| Ndufa12 | 34.45 | 18.52 | 19.29 |
| Mettl1 | 19.93 | 12.72 | 14.41 |
| Ddit3 | 43.54 | 20.69 | 23.06 |
| Gpr182 | 10.03 | 3.57 | 9.23 |
| Sec61g | 73.18 | 34.24 | 35.67 |
| Stc2 | 17.23 | 6.73 | 17.09 |
| Hint1 | 194.96 | 112.61 | 124.97 |
| Grap | 3.66 | 1.49 | 3.37 |
| 2410006H16Rik | 43.03 | 17.86 | 29.13 |
| Tvp23b | 40.04 | 27.48 | 30.46 |
| Gas7 | 20.99 | 8.25 | 9.69 |

#### Supplementary Table S2 (continued)

| NAME | Δ90+YAP$^{S127A}$ | Δ90+YAP$^{S127A}$ +NFE2L2-WT | Δ90+YAP$^{S127A}$ +L30P/R34P |
|---|---|---|---|
| Arhgef15 | 6.76 | 2.95 | 4.81 |
| Pfas | 18.64 | 12.21 | 15.19 |
| Naa38 | 18.11 | 10.14 | 10.62 |
| Tmem88 | 2.23 | 0.83 | 2.03 |
| Fam64a | 3.83 | 0.67 | 1.96 |
| Fam101b | 8.24 | 3.54 | 5.36 |
| AI662270 | 7.92 | 2.68 | 7.04 |
| Tbx2 | 9.54 | 2.02 | 3.34 |
| Ngfr | 5.83 | 1.41 | 3.12 |
| Gngt2 | 7.28 | 2.53 | 5.45 |
| Krt10 | 2.52 | 0.76 | 0.88 |
| Krt19 | 10.05 | 2.27 | 5.82 |
| Rpl27 | 320.62 | 172.54 | 199.99 |
| Icam2 | 3.31 | 1.51 | 3.22 |
| Rpl38 | 157.12 | 57.26 | 69.11 |
| Ten1 | 3.61 | 1.65 | 3.28 |
| Cygb | 44.27 | 12.66 | 14.11 |
| Mxra7 | 8.21 | 2.39 | 3.34 |
| Alyref | 45.37 | 18.49 | 21.45 |
| Colec11 | 25.86 | 7.75 | 19.31 |
| Pxdn | 14.22 | 5.24 | 10.94 |
| Gm5785 | 6.67 | 3.61 | 4.57 |
| Psma6 | 97.36 | 67.88 | 68.85 |
| Nfkbia | 48.28 | 32.95 | 36.77 |
| Rps29 | 229.39 | 47.82 | 59.9 |
| Lrr1 | 1.53 | 0.49 | 0.98 |
| Rpl36al | 209.73 | 134.15 | 136.87 |
| Rhoj | 4.05 | 1.54 | 2.61 |
| Churc1 | 19.16 | 8.47 | 10.78 |
| Vti1b | 58.5 | 41.25 | 41.74 |
| Acyp1 | 5.33 | 2.44 | 3.3 |
| Tgfb3 | 18.6 | 5.39 | 11.27 |
| Vash1 | 21.62 | 6.62 | 9.67 |
| Slirp | 19.94 | 8.9 | 9.14 |
| Ifi27l2a | 12.35 | 4.44 | 11.47 |
| Ckb | 12.03 | 5.67 | 10.47 |
| 2010107E04Rik | 53.69 | 26.47 | 28.2 |
| Stard3nl | 23.08 | 16.12 | 18.83 |
| Gmnn | 13.43 | 7.86 | 11.58 |
| Mylip | 7.91 | 3.27 | 7.56 |
| S1pr3 | 4.22 | 1.41 | 2.75 |
| Mxd3 | 7.5 | 2.63 | 3.95 |
| Tmed9 | 168.6 | 89.3 | 94.07 |
| Pcbd2 | 20.32 | 9.28 | 10.65 |
| Hnrnpa0 | 228.73 | 137.88 | 150.02 |
| Cox7c | 135.61 | 52.75 | 57.02 |
| Rps23 | 402.08 | 162.3 | 212.99 |
| Tbca | 77.78 | 34.32 | 36.66 |
| Ndufaf2 | 10.42 | 5.72 | 5.86 |
| Plpp1 | 9.61 | 5.26 | 8.92 |
| Ube2e2 | 35.83 | 23.99 | 28.01 |
| Nt5dc2 | 13.04 | 6.57 | 12.49 |
| Fam25c | 72.93 | 14.99 | 16.69 |
| Psmb5 | 34.91 | 19.56 | 26.8 |
| Mphosph8 | 24.93 | 16.02 | 21.86 |
| Mrpl57 | 25.34 | 14.51 | 16.95 |
| Pdlim2 | 4.65 | 1.26 | 2.96 |
| Zc3h13 | 36.9 | 23.65 | 30.1 |
| Commd6 | 27.61 | 17.72 | 18.54 |
| Rpl37 | 244.48 | 108.5 | 121.33 |
| Sub1 | 66.83 | 41.44 | 43.42 |
| Rpl30 | 39.53 | 19.2 | 21.45 |
| Cox6c | 108.64 | 66.95 | 67.24 |
| Eny2 | 66.04 | 46.31 | 47.52 |
| Col14a1 | 43.94 | 15.66 | 37.43 |
| Ptp4a3 | 24.09 | 10.5 | 15.69 |
| Gpihbp1 | 14.75 | 5.04 | 11.38 |
| Scx | 4.93 | 0.48 | 0.57 |
| Cyth4 | 30.5 | 13.49 | 29.85 |
| Mapk12 | 3.26 | 1.24 | 1.85 |
| Shank3 | 9.77 | 4.66 | 5.29 |
| Ccnt1 | 12.44 | 7.47 | 8.91 |
| Hnrnpa1 | 195.21 | 131.21 | 176.8 |
| Sdf2l1 | 70.64 | 32.09 | 34.75 |
| Cldn5 | 10.31 | 2.58 | 5.77 |

#### Supplementary Table S2 (continued)

| NAME | Δ90+YAP$^{S127A}$ | Δ90+YAP$^{S127A}$ +NFE2L2-WT | Δ90+YAP$^{S127A}$ +L30P/R34P |
|---|---|---|---|
| Rpl35a | 44.39 | 13.48 | 14.24 |
| Fstl1 | 25.39 | 8.6 | 13.82 |
| Cox17 | 31.4 | 15.26 | 15.74 |
| Ccdc80 | 40.64 | 10.85 | 17.4 |
| Rpl24 | 344.45 | 222.2 | 233.83 |
| Cldn8 | 2.73 | 0.05 | 0.06 |
| Erg | 6.01 | 2.56 | 4.79 |
| Tmem181a | 6.28 | 2.8 | 5.23 |
| Sft2d1 | 25.55 | 16.08 | 19.21 |
| Neurl1b | 17.68 | 8.13 | 11.18 |
| Kifc5b | 5.83 | 2.39 | 3.64 |
| Uqcc2 | 28.71 | 11.99 | 12.03 |
| Lemd2 | 57.46 | 32.21 | 35.14 |
| Mtch1 | 129.13 | 85.6 | 93.44 |
| Fgd2 | 13.39 | 5.63 | 13.14 |
| Kank3 | 8.05 | 4.28 | 4.87 |
| Rps28 | 190.68 | 58.36 | 60.68 |
| Ndufa7 | 63.75 | 41.62 | 42.33 |
| Pfdn6 | 34.17 | 19.77 | 22.13 |
| H2-Eb1 | 5.81 | 1.79 | 2.66 |
| Mrpl14 | 64.81 | 33.85 | 44.69 |
| Mea1 | 55.16 | 33.77 | 34.43 |
| 2410015M20Rik | 71.93 | 27.65 | 31.33 |
| Alkbh7 | 12.28 | 6.45 | 6.87 |
| Rab31 | 17.21 | 9.68 | 15.7 |
| Ralbp1 | 92.2 | 65.57 | 72.13 |
| Trmt61b | 5.41 | 2.86 | 3.02 |
| Lbh | 24.12 | 8.97 | 17.57 |
| Ehd3 | 23.97 | 7.55 | 15.99 |
| Cebpzos | 13.39 | 8.75 | 9.85 |
| Zmat2 | 47.98 | 34.03 | 36.01 |
| Arap3 | 14.32 | 6.9 | 10.37 |
| Pcdh12 | 6.53 | 2.76 | 4.33 |
| Tnfaip8 | 7.4 | 3.52 | 6.71 |
| Ppic | 11.71 | 4.99 | 9.68 |
| Tcf4 | 35.85 | 15.64 | 17.6 |
| Ier3ip1 | 26.89 | 17.36 | 18.16 |
| Banf1 | 56.49 | 34.36 | 36.46 |
| Drap1 | 96.56 | 41.58 | 46.89 |
| Sipa1 | 37.44 | 23.83 | 24.9 |
| Fau | 444.8 | 232.19 | 249.76 |
| Ccdc88b | 13.39 | 6.08 | 10.37 |
| Prdx5 | 183.54 | 120.57 | 134.32 |
| Polr2g | 25.72 | 16.9 | 17.48 |
| Tmem258 | 48.26 | 17.29 | 18.73 |
| Gnaq | 39.15 | 28.43 | 35.64 |
| Add3 | 24.69 | 13.06 | 15.35 |
| Ndufb11 | 115.47 | 59.53 | 65.66 |
| Ndufa1 | 35.39 | 12.82 | 13.7 |
| Ssr4 | 101.49 | 65.24 | 81.01 |
| Lage3 | 23.19 | 15.43 | 15.7 |
| Las1l | 43.24 | 31.93 | 39.36 |
| Cox7b | 75.35 | 49.31 | 58.99 |
| Rpl36a | 190.44 | 131.2 | 148.77 |
| Armcx4 | 12.17 | 2.12 | 6.48 |
| Armcx2 | 4.54 | 1.45 | 3.18 |
| Ngfrap1 | 20.58 | 11.5 | 14.1 |
| Psmd10 | 22.64 | 16.24 | 19.61 |
| Mageh1 | 2.89 | 1.4 | 2.49 |
| Eif1ax | 44.28 | 30.61 | 38.27 |
| Itih5 | 13.77 | 37.45 | 5.83 |
| A530020G20Rik | 2.73 | 6.08 | 2.33 |
| Ppbp | 3.32 | 18.82 | 3.01 |
| Zfp605 | 10.52 | 16.28 | 10.41 |
| Atoh8 | 26.18 | 57.85 | 23.54 |
| Anpep | 148.16 | 213.13 | 144.83 |
| Zfp868 | 17.01 | 25.24 | 15.3 |
| Elovl2 | 222.37 | 466.57 | 194.98 |
| Phf11d | 7.58 | 13.98 | 6.71 |
| Thpo | 16.45 | 25.23 | 14.93 |
| Lims2 | 77.18 | 126.74 | 66.89 |

**Supplementary Table S3: Previously identified *NFE2L2* point mutations in primary human HBs and HB cell lines**

| Number of tumors | NFE2L2 mutations identified | Reference |
|---|---|---|
| 24 | None | Hooks et al (6) |
| 34 | R34Gx2, D29N | Sumazin et al (7) |
| 47 + 4 cell lines | L30P, R34G, R34P, T80Ax2 | Eichenmuller (8) |
| 32 | None | Valanejad (9) |
| 53 | D77Y | COSMIC |

#### Supplementary Table S4: 46 NFE2L2 target gene transcripts from IPA

| Symbol | Synonym(s) | Entrez Gene Name | Location | Family | Entrez Gene ID | | |
|---|---|---|---|---|---|---|---|
| | | | | | Human | Mouse | Rat |
| ABCC3 | 1700019L09Rik, ABC31, ATP binding cassette subfamily C member 3, ATP-binding cassette C3, ATP-binding cassette, sub-family C (CFTR/MRP), member 3, cMOAT2, EST90757, MLP2, MOAT-D, MRP3, Multidrug Resistant Protein 3 | ATP binding cassette subfamily C member 3 | Plasma Membrane | transporter | 8714 | 76408 | 140668 |
| ATF4 | activating transcription factor 4, C/ATF, CREB-2, TAXREB67, TXREB | activating transcription factor 4 | Nucleus | transcription regulator | 468 | 11911 | 79255 |
| BRCA1 | BRCA1 DNA repair associated, BRCA1, DNA repair associated, BRCAI, BRCC1, breast cancer 1, early onset, BROVCA1, FANCS, PNCA4, PPP1R53, PSCP, RNF53 | BRCA1 DNA repair associated | Nucleus | transcription regulator | 672 | 12189 | 497672 |
| CAT | 2210418N07, ACATALASIA, Cas-1, Cat01, Catalase, Catalase1, Catl, CS1 | catalase | Cytoplasm | enzyme | 847 | 12359 | 24248 |
| CCN2 | AMPHIROGULIN, cellular communication network factor 1, cellular communication network factor 2, CTGF, CTGF isoform 1, CTGRP, Fibroblast-inducible secreted, Fisp12, HCS24, IGFBP8, IGFBP-RP2, NOV2, Tissue growth factor | cellular communication network factor 2 | Extracellular Space | growth factor | 1490 | 14219 | 64032 |
| CDH1 | AA960649, ARC-1, BCDS1, cadherin 1, Cadherin E, CD324, CDHE, CSEIL, E-cadherin, ECAD, L-CAM, Um, UVO, uvomorulin | cadherin 1 | Plasma Membrane | other | 999 | 12550 | 83502 |
| COX4I1 | AL024441, CcO IVi1, COX, COX IV-1, COX4, COX4-1, COX4A, COX4I, COXIV, cytochrome c oxidase subunit 4I1, IV-1 | cytochrome c oxidase subunit 4I1 | Cytoplasm | enzyme | 1327 | 12857 | 29445 |
| CS | 2610511A05Rik, 9030605P22Rik, Ahl4, BB234005, Cis, citrate synthase | citrate synthase | Cytoplasm | enzyme | 1431 | 12974 | 170587 |
| CXCL8 | C-X-C motif chemokine ligand 8, GCP-1, IL8, LECT, LUCT, LYNAP, MDNCF, MONAP, Monocyte-derived neutrophil chemotactic factor, NAF, NAP-1 | C-X-C motif chemokine ligand 8 | Extracellular Space | cytokine | 3576 | | |
| DDIT3 | AC144852.1, AltDDIT3, C/EBP homology, C/EBP-homologous, C/EBPzeta, Cebp Zeta, Cebp ζ, CEBPZ, CHOP, CHOP-10, DNA DAMAGE-INDUCIBLE transcript, DNA-damage inducible transcript 3, GADD153, RM4 | DNA damage inducible transcript 3 | Nucleus | transcription regulator | 1649 | 13198 | 29467 |
| G6PD | G28A, G6PD1, G6PDX, Glucose-6-P Dehydrogenase, glucose-6-phosphate dehydrogenase, glucose-6-phosphate dehydrogenase X-linked, Gpdx | glucose-6-phosphate dehydrogenase | Cytoplasm | enzyme | 2539 | 14381 | 24377 |
| GCLC | D9Wsu168e, gamma GCS HEAVY CHAIN, Gamma Glutamyl Cysteine Synthetase Light Subunit, Gamma Glutamylcysteine Synthetase, Gamma glutamylcysteine synthetase heavy subunit, GCL, GCS, GCS, Catalytic, GCS-HS, Ggcs-hs, GLCL, GLCL-H, GLCLC, glutamate-cysteine ligase catalytic subunit, Glutamate-Cysteine Ligase, Catalytic Subunit, γ Gcs, γ GCS HEAVY CHAIN, γ Glutamyl Cysteine Synthetase Light Subunit, γ Glutamylcysteine Synthetase, γ glutamylcysteine synthetase heavy subunit γ-Gcsh | glutamate-cysteine ligase catalytic subunit | Cytoplasm | enzyme | 2729 | 14629 | 25283 |
| GCLM | AI649393, Gamma gclm, gamma GCS LIGHT CHAIN, Gamma glutamylcysteine synthase (regulatory), gamma GLUTAMYLCYSTEINE SYNTHETASE, gamma-glutamylcysteine synthetase light (regulatory) subunit, Gcmc, Gcs, Regulatory, Gcs-ls, GLCLR, glutamat-cystein ligase, regulatory subunit, glutamate-cysteine ligase modifier subunit, Glutamate-Cysteine Ligase, Modifier Subunit, γ gclm, γ GCS LIGHT CHAIN, γ glutamylcysteine synthase (regulatory), γ GLUTAMYLCYSTEINE SYNTHETASE, γ-glutamylcysteine synthetase light (regulatory) subunit | glutamate-cysteine ligase modifier subunit | Cytoplasm | enzyme | 2730 | 14630 | 29739 |
| GPX2 | GI-GPx, glutathione peroxidase 2, GPRP, GPRP-2, GPX-GI, GSHPx-2, GSHPX-GI | glutathione peroxidase 2 | Cytoplasm | enzyme | 2877 | 14776 | 29326 |
| HIPK2 | 1110014O20RIK, B230339E18RIK, homeodomain interacting protein kinase 2, LOC100505582, LOC653052, PRO0593, Stank | homeodomain interacting protein kinase 2 | Nucleus | kinase | 28996 | 15258 | 362342 |
| HMOX1 | bK286B10, D8Wsu38e, haemox, HEME OXYGENASE, HEME OXYGENASE (DECYCLIZING) 1, Heme oxygenase 1, Hemox, Heox, HEOXG, Hmox, HMOX1D, HO-1, HSP32 | heme oxygenase 1 | Cytoplasm | enzyme | 3162 | 15368 | 24451 |
| IL36G | IL-1F9, IL-1H1, IL-1RP2, IL1E, interleukin 1 family, member 9, interleukin 36 gamma, interleukin 36 γ, interleukin 36, gamma, interleukin 36, γ, RGD1563019 | interleukin 36 gamma | Extracellular Space | cytokine | 56300 | 215257 | 499744 |
| ME1 | BRCAME, D9Ertd267e, HUMNDME, Malate Nadp Oxyreductase, Malic enzyme, malic enzyme 1, malic enzyme 1, NADP(+)-dependent, cytosolic, Mdh-1, MES, MOD1 | malic enzyme 1 | Cytoplasm | enzyme | 4199 | 17436 | 24552 |
| NQO1 | AV001255, DHQU, DIA4, DT-diaphorase, DTD, NAD DT-diaphorase, NAD(P)H dehydrogenase, quinone 1, NAD(P)H quinone dehydrogenase 1, NAD(P)H:quinone oxidoreductase, Nadph dehydrogenase, Nadph diaphorase, Nadph Quinone Oxidoreductase-1, NMO1, NMOR, NMOR1, NMORI, Nqo, Ox-1, Qr, QR1, Quinone reductase | NAD(P)H quinone dehydrogenase 1 | Cytoplasm | enzyme | 1728 | 18104 | 24314 |
| NR0B1 | AHC, AHCH, AHX, DAX-1, DSS, GTD, HHG, NROB1, nuclear receptor subfamily 0 group B member 1, nuclear receptor subfamily 0, group B, member 1, SRXY2 | nuclear receptor subfamily 0 group B member 1 | Nucleus | ligand-dependent nuclear receptor | 190 | 11614 | 58850 |
| OSGIN1 | 1700012B18Rik, BDGI, OKL38, oxidative stress induced growth inhibitor 1 | oxidative stress induced growth inhibitor 1 | Other | growth factor | 29948 | 71839 | 171493 |
| PGD | 0610042A05Rik, 6PGD, 6PGDH, AU019875, C78335, Cc2-27, LOC100363662, phosphogluconate dehydrogenase | phosphogluconate dehydrogenase | Cytoplasm | enzyme | 5226 | 110208 | 100360180 |
| PHGDH | 3-PGDH, 3-phosphoglycerate dehydrogenase, 4930479N23, A10, HEL-S-113, NLS, NLS1, PDG, PGAD, PGD, PGDH, PGDH3, PHGDHD, phosphoglycerate dehydrogenase, SERA | phosphoglycerate dehydrogenase | Cytoplasm | enzyme | 26227 | 236539 | 58835 |
| POMP | 2510048O06Rik, C13orf12, HSPC014, LOC100911238, PNAS-110, PRAAS2, proteasome maturation protein, proteasome maturation protein-like, RGD1305831, UMP1 | proteasome maturation protein | Nucleus | other | 51371 | 66537 | 288455<br>100911238 |
| PRDX1 | ENHANCER, Enhancer protein, Hbp23, MSP23, NKEF-A, OSF-3, PAG, PAGA, PAGB, PEROXIREDOXIN 1, peroxiredoxin 1-like 1, PEROXYREDOXIN 1, Prdx1l1, PrdxI, PRX1, PRXI, TDPX2, TDX2, TPxA | peroxiredoxin 1 | Cytoplasm | enzyme | 5052 | 18477 | 117254<br>100363379 |
| PSAT1 | D8Ertd814e, EPIP, NLS2, Phosphoserine Aminotransferase, phosphoserine aminotransferase 1, PSA, PSA1, PSAT, PSATD, Similar to phosphoserine aminotransferase | phosphoserine aminotransferase 1 | Cytoplasm | enzyme | 29968 | 107272 | 293820 |
| PSMA4 | 20S PROTEASOME alpha 4 subunit, 20S PROTEASOME α 4 subunit, C9, HC9, HsT17706, Macropain subunit C9, proteasome (prosome, macropain) subunit, alpha type 4, proteasome (prosome, macropain) subunit, α type 4, proteasome subunit alpha 4, Proteasome subunit c9, proteasome subunit α 4, Proteosome subunit α 4, PSC9 | proteasome subunit alpha 4 | Cytoplasm | peptidase | 5685 | 26441 | 29671 |
| PSMA5 | Aa409047, Macropain zeta chain, Macropain ζ chain, proteasome (prosome, macropain) subunit, alpha type 5, proteasome (prosome, macropain) subunit, α type 5, proteasome subunit alpha 5, proteasome subunit α 5, PSC5, ZETA, ζ | proteasome subunit alpha 5 | Cytoplasm | peptidase | 5686 | 26442 | 29672 |

### Supplementary Table S4: 46 NFE2L2 target gene transcripts from IPA( continue)

| Symbol | Synonym(s) | Entrez Gene Name | Location | Family | Entrez Gene ID | | |
|---|---|---|---|---|---|---|---|
| | | | | | Human | Mouse | Rat |
| PSMB2 | AU045357, AW108089, beta 2 PROTEASOME subunit, C7-I, D4Wsu33e, HC7-I, proteasome (prosome, macropain) subunit, beta type 2, proteasome (prosome, macropain) subunit, β type 2, proteasome subunit beta 2, proteasome subunit β 2, β 2 PROTEASOME subunit | proteasome subunit beta 2 | Cytoplasm | peptidase | 5690 | 26445 | 29675 |
| PSMB5 | 26s Proteasome Beta 5, 26s Proteasome β 5, beta 5 PROTEASOME subunit, Lmp17, LMPX, MB1, proteasome (prosome, macropain) subunit, beta type 5, proteasome (prosome, macropain) subunit, β type 5, Proteasome 20S X, proteasome subunit beta 5, proteasome subunit β 5, X proteasome subunit, β 5 PROTEASOME subunit | proteasome subunit beta 5 | Cytoplasm | peptidase | 5693 | 19173 | 29425 |
| PSMD4 | AF, AF-1, angiocidin, ASF, MCB1, proteasome (prosome, macropain) 26S subunit, non-ATPase, 4, Proteasome 26s subunit non-atpase 4, proteasome 26S subunit, non-ATPase 4, pUB-R5, Rpn10, S5A | proteasome 26S subunit, non-ATPase 4 | Cytoplasm | other | 5710 | 19185 | 83499 |
| S100P | MIG9, PLACENTAL CALCIUM binding, S100 calcium binding protein P | S100 calcium binding protein P | Cytoplasm | other | 6286 | | |
| SERPINE1 | beta MIGRATING PLAS ACTIVATOR, beta-MIGRATING PLASMINOGEN ACTIVATOR INHIBITOR I, PAI, PAI-1, PAI1A, Pai1aa, Planh, PLANH1, Plasminogen activator inhibitor 1, RATPAI1A, serine (or cysteine) peptidase inhibitor, clade E, member 1, serpin family E member 1, SERPINE, β MIGRATING PLAS ACTIVATOR, β-MIGRATING PLASMINOGEN ACTIVATOR INHIBITOR I | serpin family E member 1 | Extracellular Space | other | 5054 | 18787 | 24617 |
| SHC1 | p52SHC, p66, P66shc, SHC, Shc (46 kDa isoform), SHC adaptor protein 1, Shc p66 isoform, SHCA, src homology 2 domain-containing transforming protein C1 | SHC adaptor protein 1 | Cytoplasm | other | 6464 | 20416 | 85385 |
| SHMT2 | 2700043D08Rik, AA408223, AA986903, GLYA, HEL-S-51e, serine hydroxymethyltransferase 2 (mitochondrial), Serine Hydroxymethyltransferase2, SHMT | serine hydroxymethyltransferase 2 | Cytoplasm | enzyme | 6472 | 108037 | 299857 |
| SLC7A11 | 9930009M05RIK, AI451155, CCBR1, CYSTEINE GLUATAMINE TRANSPORTER, solute carrier family 7 (cationic amino acid transporter, y+ system), member 11, solute carrier family 7 member 11, sut, xCT | solute carrier family 7 member 11 | Plasma Membrane | transporter | 23657 | 26570 | 310392 |
| SNAI2 | SLUG, SLUGH, SLUGH1, snail family transcriptional repressor 2, snail family zinc finger 2, SNAIL2, WS2D | snail family transcriptional repressor 2 | Nucleus | transcription regulator | 6591 | 20583 | 25554 |
| SOD1 | ALS, ALS1, B430204E11Rik, Cu/Zn-SOD, CuZnSOD, czSOD, HEL-S-44, hSod1, Ipo-1, IPOA, SOD, SOD1L1, SODC, superoxide dismutase 1, superoxide dismutase 1, soluble | superoxide dismutase 1 | Cytoplasm | enzyme | 6647 | 20655 | 24786 |
| SOD2 | IMAGE:4711494, IPO-B, MANGANESE DEPENDENT SOD, Manganese Superoxide Dismutase, Manganese Superoxide Dismutase 2, MGC5618, MITOCHONDRIAL SOD, Mn superoxide dismutase, MNSOD, mtSOD, MVCD6, superoxide dismutase 2, superoxide dismutase 2, mitochondrial | superoxide dismutase 2 | Cytoplasm | enzyme | 6648 | 20656 | 24787 |
| SRGN | haematoPOETIC PROTEOGLYCAN CORE, HEMATOPOETIC PROTEOGLYCAN CORE, PGSG, PPG, PRG, PRG1, Serglycin, Sgc | serglycin | Cytoplasm | other | 5552 | | |
| TALDO1 | TAL, TAL-H, TALDOR, Transaldolase, transaldolase 1 | transaldolase 1 | Cytoplasm | enzyme | 6888 | 21351 | 83688 |
| TFAM | AI661103, Hmgts, MTDPS15, MTTF1, MTTFA, TCF6, TCF6L1, TCF6L2, TCF6L3, Tfa, transcription factor A, mitochondrial, tsHMG | transcription factor A, mitochondrial | Cytoplasm | transcription regulator | 7019 | 21780 | 83474 |
| TKT | HEL-S-48, HEL107, p68, SDDHD, TK, TKT1, TRANSKETOLASE | transketolase | Cytoplasm | enzyme | 7086 | 21881 | 64524 |
| UGT1A1 | BILIQTL1, BILIRUBIN UDP-GLUCURONOSYLTRANSFERASE ISOENZYME1, GNT1, HUG-BR1, UDP glucuronosyltransferase 1 family, polypeptide A1, UDP glucuronosyltransferase family 1 member A1, Udp Glycosyltransferase 1, UDPGT, UDPGT 1-1, Udpgt-1a, UGT1, UGT1A, UGT1A01, Ugt1a2b, UgtBr1 | UDP glucuronosyltransferase family 1 member A | Cytoplasm | enzyme | 54658 | 394436 | 24861 |
| UGT1A7 (incl | A13, GNT1, HLUGP4, hUG-BR1, LOC100048744, LUGP4, mUGTBr/P, UDP glucuronosyltransferase 1 family, polypeptide A9, UDP glucuronosyltransferase family 1 member A10, UDP glucuronosyltransferase family 1 member A7, UDP glucuronosyltransferase family 1 member A8, UDP glucuronosyltransferase family 1 member A9, UDP glycosyltransferase 1 family, polypeptide A10, UDPGT, UDPGT 1-7, UDPGT 1-8, UDPGT 1-9, UGT-1A, UGT-1G, UGT-1H, UGT-1I, UGT-1J, UGT1, UGT1-01, UGT1-07, UGT1-08, UGT1-09, UGT1-10, UGT1-9, UGT1.1, UGT1.10, UGT1.8, UGT1.9, UGT1A1, UGT1A10, Ugt1a11, Ugt1a12, Ugt1a13, UGT1A8, UGT1A8S, UGT1A9, UGT1A9S, UGT1AI, UGTP4 | UDP glucuronosyltransferase family 1 member A | Cytoplasm | enzyme | 54577<br>54576<br>54575<br>54600 | 394434<br>394430 | 301595 |

**Supplementary Table S5. Antibodies used in the current study**

| NAME OF ANTIBODY | VENDOR | CATALOG NO. | DILUTION USED |
|---|---|---|---|
| **CPT1A** | Abcam | ab128568 | 1:1000 |
| **GAPDH** | Sigma | G8795 | 1:10,000 |
| **GLUT1** | Abcam | ab115730 | 1:20,000 |
| **GLUTAMINE SYNTHETASE (GLNS)** | GeneTex | GTX109121 | 1:5000 |
| **GLUTAMINASE (GLS)** | Abcam | ab131554 | 1:1000 |
| **β-CATENIN** | Abcam | ab16051 | 1:4000 |
| **PAI1(SERPINE1)** | R&D | AF3828sp | 1:1000 |
| **PDHA1** | Santa Cruz | SC-377092 | 1:1000 |
| **$_P$PDHA1 (PSER$^{293}$)** | Calbiochem | AP1062 | 1:1000 |
| **PC** | Abcam | Ab128952 | 1:2000 |
| **PKM1** | CST | 7067 | 1:1000 |
| **PKM2** | CST | 3198 | 1:1000 |
| **PFK-L** | Aviva System Biology | ARP45774 | 1:1000 |
| **GLUT2** | ProteinTech | 20436-1-AP | 1:300 |
| **GLUT4** | CST | 2213 | 1:1000 |
| **HISTONE H3** | CST | 9715 | 1:2000 |
| **GLUD1** | CST | 12793 | 1:1000 |
